# Enhanced Calcium Signaling in BLA Pyramidal Neurons Underlies a Sex and Circuit Specific Amygdala Dysfunction After Global Cerebral Ischemia

**DOI:** 10.1101/2025.01.28.635307

**Authors:** Jose J. Vigil, Erika Tiemeier, Nicholas E. Chalmers, Paco S. Herson, Nidia Quillinan

**Author notes:** **Corresponding Authors:** Nidia Quillinan, PhD Associate Professor, Department of Anesthesiology, University of Colorado School of Medicine, Anschutz Medical Campus 12801 E. 17^th^ Ave Aurora, CO 80045, Paco S. Herson, PhD, Professor and Associate Dean for Research Innovation, Department of Neurological Surgery, The Ohio State University College of Medicine, 460 W. 12^th^ Ave. Columbus, OH 43210. Equal contributing authors (Paco S. Herson, Nidia Quillinan).

## Abstract

While advances in resuscitation science have improved cardiac arrest survival, we lack therapies to improve cognitive-affective outcomes in this patient population. Our lab has previously identified cognitive dysfunction in a mouse model of global cerebral ischemia (GCI) which has been attributed to hippocampal neurodegeneration and impaired hippocampal plasticity. However, no study has attempted to identify amygdala dysfunction after GCI, despite clinical evidence of emotional dysfunction, such as anxiety and Post-Traumatic Stress Disorder (PTSD). Therefore, it is important to identify the effect that GCI has on the amygdala, the emotional center of the brain. Our lab has a well-developed, translatable mouse model of GCI, the cardiac arrest/cardiopulmonary resuscitation model (CA/CPR), that has been instrumental in assessing amygdala function after GCI. We have utilized the amygdala-dependent delay-fear conditioning (DFC) paradigm to assess associative learning and memory and have performed field excitatory post-synaptic potential (fEPSP) recordings in two circuits within the amygdala, as measures of amygdala function. We have found a sex- and circuit-specific deficit in LTP of the cortical input to the basolateral amygdala (BLA), after GCI, that corresponds with a male specific associative learning and memory deficit. We found no evidence that these deficits of amygdala function can be attributed to GCI-induced neurodegeneration within the amygdala or altered locomotor function. We did, however, find that neuronal dendritic spine calcium signaling dynamics are enhanced in male survivors of GCI and are likely contributing to the observed LTP deficit and ultimately the deficit in associative learning and memory processes.

**Significance Statement:** While cognitive dysfunction has been well characterized in CA/CPR survivors, the development of emotional dysfunction that may arise in survivors is understudied. Poor follow up of CA/CPR survivors has left a gap in our knowledge about the effect that GCI has on the amygdala, the emotional center of the brain. Thus, this study utilized a pre-clinical mouse model of CA/CPR to characterize amygdala dysfunction that arises from GCI. We found a male-specific deficit in associative learning and memory that corresponds with a circuit specific LTP deficit, which arises from elevated calcium signaling in dendritic spines. Thus, this study provides valuable information and rationale for extended follow up of CA/CPR survivors to identify and treat the development of GCI-induced affective disorders.

## Introduction

Cardiac arrest (CA) is one of the leading causes of death and disability in humans. The estimated global incidence of out-of-hospital cardiac arrest is 55 per 100,000 people per year (Yan et al., 2020). Mortality from CA remains high and resuscitated survivors experience neurological impairments that negatively impact quality of life. Cognitive deficits are a well-documented long-term consequence of global cerebral ischemia (GCI) induced by CA (Chew et al., 2018; Cohan et al., 2015; Elmer & Callaway, 2017) and it is becoming apparent that CA survivors also experience severe emotional dysfunction. An estimated 40-60% of CA survivors experience depression and anxiety and 30% experience Post-Traumatic Stress Disorder (PTSD) (Naber & Bullinger, 2018). Development of these affective disorders in survivors of CA suggest amygdala dysfunction as an underlying cause. Yet amygdala dysfunction resulting from CA is not well characterized in clinical studies or animal models. Therefore, this study fills an important gap in knowledge by using a translational mouse model of GCI, the Cardiac Arrest/Cardiopulmonary Resuscitation (CA/CPR) mouse model, to explore deficits in amygdala function that may contribute to emotional dysfunction after GCI.

A major function of the amygdala is to mediate associative learning and memory processes, the ability to learn and remember relationships between different stimuli(Krabbe et al., 2018; Maren, 2005; Rogan et al., 1997). Dysfunction of associative learning and memory processes have been implicated as the underlying cause of neuropsychiatric disorders (Allen et al., 2019; Diener, 2013; Grillon, 2002), and GCI-induced changes in associative learning can be assessed with the delay-fear conditioning (DFC) paradigm(Raybuck & Lattal, 2011; Y. Sun et al., 2020). The ability to learn associative relationships during DFC is amygdala dependent and synaptic plasticity in the basal and lateral sub-nuclei of the amygdala (referred to as the BLA) are vital for associative learning(Anglada-Figueroa & Quirk, 2005; Meis et al., 2018; Phillips & LeDoux, 1992; Ressler, 2010; Rogan et al., 1997; Y. Sun et al., 2020; Tsvetkov et al., 2002). Synaptic plasticity is the activity dependent change in synaptic transmission that is the cellular mechanism of learning and memory. One form of synaptic plasticity, Long-Term potentiation (LTP), is the activity dependent sustained increase in synaptic transmission and is known to underlie amygdala dependent associative learning and memory (Mahan & Ressler, 2012; Rogan et al., 1997).

During associative learning and memory tasks such as DFC, sensory information converges onto the BLA before being distributed throughout the amygdala for processing and subsequent output to other brain regions, to mediate the appropriate behavioral response(Janak & Tye, 2015; LeDoux, 2007). LTP of the cortical inputs to the BLA require both functional NMDARs and LTCCs and is essential for associative learning and memory(Bauer et al., 2002; Tsvetkov et al., 2002). BLA neurons also make synaptic contacts with each other and elevated synaptic activity within this intra-amygdala circuit is associated with anxiety-like behavior(Rosen & Schulkin, 2022). LTP of intra-amygdala BLA synapses requires functional NMDARs, but not LTCCs(Drephal et al., 2006). Dysfunction of either ion channel will alter calcium dynamics within the neuron and influence associative learning despite their circuit specific effects on LTP. Thus, circuit specific alterations of LTP in the BLA can implicate specific calcium sources as the underlying cause of dysfunctional associative learning and memory processes after GCI.

Here, we have revealed a male-specific deficit in DFC after GCI that correlates with diminished LTP in the cortical input (LTCC- and NMDAR-dependent LTP) to the BLA but not LTP of the intra-amygdala circuitry (NMDAR-dependent LTP). The discovery of this dysfunction in associative learning and memory processes and identification of an underlying circuit specific dysfunction of LTP after GCI begins to address a gap in our knowledge about the effects of GCI on the brain. This circuit specific deficit in BLA plasticity also suggests functional changes in dendritic spine calcium dynamics as the underlying molecular mechanism for the sex and circuit specific amygdala dysfunction we have discovered.

## Materials and Methods

### Animals

All experimental protocols were approved by the Institutional Animal Care and Use Committee (IACUC) at the University of Colorado, Anschutz Medical Campus and adhered to the National Institute of Health guidelines for the care and use of animals in research. Male and female mice were included throughout the study, unless otherwise indicated. All animals were 8-12 weeks of age at the time of surgery. All mice were permitted free access to water and standard lab chow ad libitum with a 14/10-h light/dark cycles.

### Cardiac Arrest/Cardiopulmonary Resuscitation

As we have previously described(Orfila et al., 2018) young adult (8-12 weeks) Charles River C57BL/6 mice were anesthetized using 3% isoflurane and intubated with 20% oxygen (100cc/min) 80% air mixture (900cc/min). Respiration was set to 160 strokes/minute with a stroke volume of 80-150µL (calculated as eight times mouse’s body weight in grams with a maximum of 150 µL). A jugular catheter was inserted, and ECG (Corometrics Eagle 4000N., Milwaukee WI) electrodes were connected to display heart rate. Using a heat lamp and water-filled head coil, body and head and temperatures were brought to 37.5±0.5°C. To initiate cardiac arrest (CA), 0.05mL potassium chloride (0.5mEq/mL) was injected via jugular catheter, the respirator was disconnected, and asystole was confirmed for 8 minutes via ECG readout. During CA, head temperature was maintained at 37.5±0.5°C using a 40°C water-heated head coil while body temperature was permitted to fall to a minimum temperature of 35.5°C. Mechanical ventilation (MiniVent Type 845., Germany) was initiated 30 seconds prior to resuscitation at a rate of 210 breaths/minute with 100% oxygen at 300cc per minute. Resuscitation was initiated via chest compressions and epinephrine (up to 0.5mL, 16µg/mL) was administered via the jugular catheter over a period of 3 minutes until normal sinus rhythm was achieved. After recovery of spontaneous respiration rate of 30 breaths/minute, air was increased to 300cc/minute and respiration rate was reduced by 10 strokes/minute for 5 minutes until extubating. Animals were excluded if the return of spontaneous circulation was not achieved within 3 minutes of initiating compressions or if return of spontaneous breathing with successful extubation was not achieved within 30 minutes after resuscitation. Mice were single housed after recovery from anesthesia and post-surgical care consisted of 2-3 days of housing cages on a warming pad, provision of soft chow, and daily administration of 0.5mL subcutaneous 0.9% saline.

### Delay Fear Conditioning

#### Apparatus

The fear conditioning apparatus consisted of a sound attenuating chamber (24” L x 14” W x 14”H) which contained a grid-shock floor (Coulbourn, Holliston, MA) onto which a 5-sided plexiglass cube (6 x 6 x6) was placed to contain the mice during the experiments (Context A). The shock floor was connected to a shock stimulator (Coulbourn, Holliston, MA) that was automatically triggered by an Arduino control board. The conditioned stimulus (CS) was a 15s tone (60 dB) delivered by a piezo buzzer (Amazon) that co-terminated with foot-shock (0.5mA, 1s) (unconditioned stimulus – US). For all procedures carried out in Context A, the apparatus was cleaned with a 70% isopropanol solution before and after each animal, and each animal was transported to the conditioning room in their home-cage. Context B consisted of a white bucket (7” diameter, 7” tall) loosely covered by a clear plexiglass sheet and placed into the sound attenuating chamber. A small vial containing vanilla extract was placed in the corner of the sound attenuating chamber and the apparatus was cleaned with a 70% ethanol solution before and after each animal. For all procedures carried out in Context B, the animals were individually transported to the conditioning room in small, empty, white square containers. Any-maze software (Stoelting, Wood Dale, IL) was used to record and assess freezing behavior during every stage of the procedure.

#### Procedure

Seven days post-CA/CPR, adult (8-12 week) male and female mice were subjected to delay-fear conditioning to amygdala-dependent cued fear learning. Male and female animals were tested on separate weeks to avoid hormonal influence on behavioral outcomes. On the first day of the procedure, animals were habituated to the conditioning room while in their home cage for one hour. Animals were also habituated to Context A for 5 minutes without presentation of the CS or the US. 24-hours later, animals were re-exposed to Context A to obtain a 2-minute baseline before CS/US pairing. After the baseline period, three CS/US pairings were delivered with 1-minute separating each pairing. After the last CS/US pairing, mice were allowed to remain in the chamber for 1 minute and 45 seconds before being returned to their home cage. The next day, animals were returned to context A for 2-minutes without presentation of CS or US to assess contextual fear and extinguish any non-associative cues(Poon & Schmid, 2012; Smith et al., 2007). The following day, animals were transported to the conditioning room in novel containers and exposed to the novel context B for 2 minutes. After this baseline period, the CS was delivered to assess amygdala-dependent cued fear. Mice were allowed to remain in context B for an additional 1.5 minutes after the CS presentation before being returned to their home cage.

### Open Field Test

#### Apparatus

The open field (OF) apparatus consisted of a square white chamber (14” L x 14” W x 13”H) with an open top. A camera was mounted above each chamber to record the activity of the mice during the test. The chamber was divided into two zones, an inner zone was indicated with permanent marker on the chamber floor for consistent placement of measurement zones within the recording and analysis software, Any-maze (Stoelting, Wood Dale, IL). The outer zone was the rest of the chamber surrounding the inner zone. A small lamp was placed in the behavior room to ensure a consistent lighting level (∼20 lux) and to ensure there were no shadows cast into the chamber. The chamber was cleaned with 70% isopropanol between animals.

#### Procedure

Seven days post-CA/CPR, adult (8-12 weeks) mice were subjected to the OF test to assess anxiety like behavior and locomotor activity. Mice subjected to the OF test were a separate cohort from those that had undergone DFC testing. Animals were habituated to the behavior room in their home cage for 1-hour before beginning the testing procedure. At the start of testing, an animal was placed into the chamber and video recording was begun in Any-Maze (Stoelting, Wood Dale, IL). The animal was allowed to explore the apparatus for 10 minutes before being returned to its home cage. Time spent in the inner and outer zone was assessed by the software and percentage of time spent in the open (center zone) was calculated by dividing the amount of time spent in the center of the apparatus by the total time of the test and multiplying by 100. Average speed and total distance traveled was also calculated within Any-maze (Stoelting, Wood Dale, IL) for each animal.

### FluoroJade Imaging and Immunohistochemistry

#### Brain and slice preparation

Seven days post-CA/CPR, adult (8-12 week) male mice were transcardially perfused with ice cold phosphate buffered saline (PBS) for 5 minutes followed by 5 minutes of ice cold 4% paraformaldehyde (PFA). The brains were then rapidly extracted and placed in PFA overnight at 4°C. The next day, the brains were placed into cryoprotection solution (20% glycerol, 20% Sorenson’s Buffer, 60% water) at 4°C until the brain sank to the bottom of the vial (overnight). The cryoprotected brains were then stored at 4°C until slicing of the brain. Brains were sliced in the coronal orientation (50µm) on a sliding microtome (Leica SM 2010R, Germany) using OTC compound and crushed dry ice to secure the brain to the cutting stage and freeze the brain itself. Slices were collected with a small paint brush and placed into a 24-well plate containing cryostorage (50% 0.2M PO4, 30% ethylene glycol, 1% polyvinylpryyolidine, 19% sucrose) solution until processing for immunohistochemistry and FluoroJade staining.

#### IHC and FluoroJade Staining

Free floating sections were washed 3x for 5-minutes each in Phosphate Buffered Saline (PBS) before blocking in PBS-Triton X (PBST)solution containing 5% Normal Donkey Serum (NDS) for 1-hour at room temperature. Slices were then incubated in primary rabbit anti-mouse NeuN monoclonal antibody (Abcam, Waltham, MA) at 1:500 in 3% PBST solution overnight at 4°C. Slices were then washed 3x in PBS for 5-minutes each and incubated in donkey anti-rabbit (Jackson ImmunoResearch, Westgrove, PA) secondary antibody conjugated to AlexaFluor 594 at 1:500 in PBST containing 3% NDS for 1-hour at room temperature. Sections were then washed 3x in PBS for 5 minutes before mounting onto charged slides and being allowed to adhere overnight at room temperature. Slides were then successively incubated in Basic EtOH (20mL of 5% NaOH solution into 80mL absolute alcohol) for 5-minutes, 70% EtOH for 2-minutes, 50% EtOH for 2-minutes before being washed in dH_2_O for 2-minutes. Slides were then submerged in a 0.06% Potassium Permanganate blocking solution for 15-minutes and then placed into the FluoroJade B (Biosensis, Australia) working solution (4mL of 0.01% stock FJ solution (0.01g of dye powder into 100mL H_2_O) into 96mL of 0.1% acetic acid solution) for 20-minutes. Slides were then rinsed in dH_2_O 3x for 1-minute each and then allowed to dry at 60°C for 10-minutes. Slides were then cleared in Xylene for 2-minutes before being cover slipped in DPX (Sigma, St. Louis, MO) mounting media.

#### Confocal Imaging

Confocal images were acquired on an Olympus FV1000 laser scanning microscope (Olympus, Bartlett, TN) using a 20x objective and FluoView software (Olympus FluoView, FV10-ASW, Bartlett, TN). Z-stacks were acquired at a thickness of 0.75µM and obtained throughout the entire depth of the slice. Multiple fields of view were acquired to encompass the entire BLA complex and subsequently stitched together to allow visualization of the entire BLA. Two successive brain sections and both hemispheres containing the BLA from each section were imaged in each animal from Bregma −1.22mm to ∼ Bregma −1.46mm (The mouse brain in stereotaxic coordinates, Third Edition). Analysis was performed in ImageJ (NIH) using the cell counter plugin to quantify the number of NeuN^+^ or FluoroJade positive cells. Hippocampal images from the same sections were also obtained to corroborate previous results from our lab but the numbers of FJ or NeuN positive cells were not quantified. The values obtained from each section in an individual mouse were averaged to yield one value per animal, thus n = number of animals.

### Electrophysiology

#### Slice Preparation

Amygdala containing slices were prepared 7-days after recovery from CA/CPR or sham surgeries as previously reported. Mice were anesthetized with 5% isoflurane in an O_2_-enriched chamber and transcardially perfused with ice-cold (2–5 °C) oxygenated (95% O2/5% CO_2_) artificial cerebral spinal fluid (ACSF) for 2-minutes prior to decapitation. The brains were then rapidly extracted and sectioned in ice-cold ACSF. The composition of ACSF was the following (in mM): 126 NaCl, 2.5 KCl, 25 NaHCO3, 1.3 NaH2PO4, 2 CaCl2, 1 MgCl2, and 10 glucose. Transverse hippocampal slices (300μm thick) were cut with a Vibratome VT1200S (Leica., Deer Park IL) and transferred to a holding chamber containing room temperature ACSF for at least 1-hour before recording. For 2-Photon imaging experiments, it was necessary to optimize the cutting procedure to ensure the availability of more “near surface” neurons and increase the odds of successful experiments. This was done by perfusion and sectioning in a modified ACSF containing (in mM): 126 NaCl, 2.5 KCl, 25 NaH2CO3, 1.3 NaH2PO4, 1 CaCl2, 3 MgSO4, and 10 glucose. After sectioning, these sections were transferred to a holding chamber containing this modified ACSF at 34°C for 30 minutes before being transferred to a holding chamber containing normal ACSF at room temperature for at least 30 minutes prior to recording.

#### Field Electrophysiology

Synaptically evoked field potentials were recorded from slices containing the BLA, which were placed on a temperature controlled (31 ± 0.5 °C) submersion chamber perfused with ACSF at a rate of 2.5 mL/min. Field excitatory post-synaptic potentials (fEPSP) were produced by delivering current pulses (ISO Flex Stimulus isolation unit, A.M.P.I, Israel) to either the external capsule (cortical input to the BLA) and recording within the BLA or by stimulating within the anterior part of the BLA and recording within the posterior part of the BLA (intra-amygdala). Analog fEPSPs were amplified (1000×) and filtered through a preamplifier (Model LP511 AC, Grass Instruments., West Warwick, RI) at 1.0 kHz, digitized at 10 kHz and stored on a computer for later off-line analysis (Clampfit 10.4, Axon Instruments, Union City, CA). The peak response of the fEPSP was measured by delivering test pulses of increasing stimulus intensity every 20 s in 50µA increments (0-500µA) to obtain an input/output (I/O) curve. The voltage response was adjusted to 50% of the maximum peak amplitude derived from the I/O curve, and test pulses were evoked every 20 s. Paired-pulse responses were recorded using a 50-ms inter-pulse interval (20 Hz) and expressed as a ratio of the peak amplitude of the second pulse over the peak amplitude of the first pulse. A 20-min stable baseline was established before delivering four theta burst stimulations (TBS- train of four pulses delivered at 100 Hz in 30-ms bursts repeated 10 times with 200-ms inter-burst intervals) at 20 second intervals. Following TBS, the fEPSP was recorded for an additional 60 min. The average 10-min peak amplitude from 50 to 60 min after TBS was divided by the average of the last 10-min of the baseline (set to 100%) prior to TBS, to determine the magnitude of potentiation. For time course graphs, normalized fEPSP peak amplitude values were averaged and plotted as the percentage of baseline. A maximum of two slices per animal were used for analysis and averaged per animal (n = number of animals).

#### Whole-Cell Patch Clamp Electrophysiology

After recovery, brain slices were transferred to the submersion recording chamber and superfused with ACSF (22-25°C for LTCC recordings and 31±0.5°C for 2 photon/electrophysiology recordings) at a flow rate of 2.5 ml/min. Recordings were obtained from pyramidal neurons within the BLA area visually identified with a differential interference contrast microscope (Leica DMFLFS, Germany) and an infrared CCD camera under 63x magnification. Recording electrodes made of borosilicate glass were constructed using a vertical electrode puller (Narishige PC-100, Amityville, NY) and had a resistance of 3-5 MΩ when filled with intracellular solution. Responses were amplified and filtered at 5 kHz (Multiclamp 700B, Axon Instruments, Union City, CA) and digitized at 20kHz (Digidata 1440A and Clampex 10.7, Axon Instruments, Union City, CA). No series resistance compensation or junction potential corrections were performed. Whole-cell configuration of BLA neurons was achieved at −65mV after achieving a >1GΩ seal. Access resistance was monitored throughout the experiments by delivering a 5mV hyperpolarizing step throughout the recordings. If access resistance was >30MΩ or changed by >20% during any recording, it was excluded from analysis. For some of the recordings, neurobiotin (Vector Laboratories, Newark, CA) was used in the internal solution (1 mg/mL) for anatomical visualization within the BLA.

For LTCC recordings the internal solution contained (in mM): 110.45 CsMeSO_3_, 2.5 CsCl, 8 NaCl, 7 TEA-Cl, 20 HEPES, 0.2 EGTA, 5 Mg-ATP, 0.3 Na-GTP, 5 QX-314, and neurobiotin. The ACSF solution contained AP5 (50µM), DNQX (10µM), and Gabazine (10µM) to block synaptic transmission. Upon the establishment of whole cell configuration, cells were maintained at a holding potential of −65mV in voltage clamp configuration for 10 minutes before beginning the experiment to allow for diffusion of intracellular contents. To record whole-cell calcium currents the neuron was held at −50mV and depolarizing voltage steps (500ms duration) were delivered from −50 to +40mV in 10mV increments. Five of these voltage step recordings were obtained before beginning perfusion of Nifedipine (10µM) containing ACSF. After 10 minutes of Nifedipine perfusion, five additional voltage clamp recordings were obtained. Current measurements were made 400ms after the onset of voltage step and the values obtained were normalized to membrane capacitance (C_m_). Normalized current measurements obtained during Nifedipine treatment were then subtracted from the values obtained prior to drug treatment, which allowed us to quantify Nifedipine sensitive current (current attributed to LTCCs). Due to the irreversible inhibition of LTCCs by Nifedipine, only one recording could be made per hemisphere, with a maximum of two recordings per animal. If more than one recording was obtained per animal, the values were averaged to yield one value per animal (n = number of animals).

For whole-cell recordings obtained with concurrent 2-Photon calcium imaging of dendritic spines, the internal solution contained (in mM): 130 Potassium Gluconate, 10 HEPES, 2 MgCl, 10 Phosphocreatine di(tris) salt, 2 Na-ATP, 0.2 Na GTP, 0.01 Alexa 594, and 0.2 Cal 520. Upon the establishment of whole cell configuration, cells were maintained at a holding potential of −65mV in voltage clamp configuration for 10 minutes before beginning the experiment to allow for allow for intracellular equilibration of the calcium indicator (Cal 520 potassium salt, AAT Bioquest, Pleasanton, CA) and cell fill (Alexafluor 594 hydrazide, Thermo Fisher Scientific, Carlsbad, CA). During this time, a glass monopolar stimulating electrode containing 150 mM NaCl and 10µM Alexafluor 594 was placed near a visually identified secondary dendritic branch ∼50-100µM away from the soma. Current pulses (0-1µA) were delivered every 20 seconds and were adjusted to yield a voltage response ∼10mV in amplitude (ISO Flex Stimulation isolation unit, A.M.P.I, Israel). Upon successful acquisition of synaptically evoked excitatory post synaptic potentials (EPSPs), 2-photon imaging parameters were set within the imaging acquisition software (PrarieView, Bruker, Billerica, MA) to be able to visualize and quantify calcium transients in distal dendritic spines (see below). First, a current clamp experiment was performed by holding the cell at −65mV and delivering depolarizing current steps (25pA) from 0-425pA and recording the voltage response and corresponding calcium transient in distal dendritic spines. Second, a stable 5-minute baseline of evoked EPSPs and calcium transients were recorded at −70mV before delivering four TBS (as in the above fEPSP recordings) at −60mV and then continuing to record the evoked voltage response for an additional 5-minutes at −70mV, while recording the corresponding calcium response in distal dendritic spines.

#### 2-Photon Calcium Imaging

Two-photon fluorescence images were obtained using a Bruker Ultima Investigator DL (Bruker, Billerica, MA) with an Olympus BX51WIF microscope (Olympus, Bartlett, TN) equipped with a Mai Tai-Deep See laser (Spectra Physics, Milpitas, CA) for two-photon excitation and 60x water immersion objective. Neurons were loaded with the green, fluorescent calcium indicator cal-520 (200 nM; Kd = 300 nM; AAT Bioquest, Pleasanton, CA) and the red fluorescent calcium-insensitive AlexaFluor 594 (10 μM; Thermo Fisher Scientific, Carlsbad, CA). Z-stacks were acquired with no optical zoom at 1024 x 1024 resolution with 3µM step size in the z-axis, with 3x optical zoom at 1024 x 1024 with 0.5µM z-step size, and with 10x optical zoom and 0.02µM z-step size. Fluorescence images were acquired in line scan-mode (1.25 MHz) at a resolution of 25.64 pixels/μm. Calcium signals were calculated as the change in green fluorescence (Cal 520) normalized to the red fluorescence (Alexa 594), G/R for each region of interest. G/R values were then normalized to the fluorescent signal before any currents steps or electrical stimulation were delivered to yield values of ΔF/F. Data collection was limited to spines on secondary dendritic branches approximately 50-100µM from the soma. Calcium signals were analyzed offline using Image J-Fiji (NIH). For current clamp experiments, the area under the curve of the calcium response in individual spines was calculated in Prism (Graph Pad, San Diego, CA) and plotted as a function of current intensity delivered to the neuron. The peak of the calcium response was also obtained from the data for recordings that contained a single action potential at rheobase for comparison. The decay of the intrinsic calcium response to a single action potential was fit with a single-phase decay exponential, and tau values were used for comparison of the decay of the calcium response. For analysis of synaptic calcium responses to TBS, only the area under the curve (Prism, Graph Pad, San Diego) of the ΔF/F plot was used for comparison between groups.

#### Rigor and Statistics

To enhance the rigor and reproducibility of our study, all experimenters were blind to injury status of animals and drug treatment throughout data acquisition. All experiments followed ARRIVE 2.0 guidelines. Data was compiled and grouped for statistical comparison after completion of all data acquisition, to limit experimenter bias. All animals were randomly assigned to experimental groups by a surgeon who performed all CA/CPR, and a separate experimenter performed all acquisition and analysis of data to ensure blinding. Sample size and power analysis were performed with a small subset of preliminary data using G-power (UCLA, Los Angeles, CA) to detect a 20% change with 90% power. All data are expressed as mean ± SD unless otherwise stated. Normality of data was assessed with the Anderson-Darling normality test(“Anderson–Darling Test,” 2008) to ensure the proper statistical tests were performed. Where appropriate, either a two-way ANOVA, Student’s T-Test, Mann-Whitney U test, or Nested ANOVA was performed to assess statistical significance. Differences were considered statistically significant if p < 0.05. Either Prism (Graph Pad, San Diego, CA) or R (R version 4.4.1, http://www.r-project.org) was used for statistical analysis.

## Results

### Associative Learning and Memory is Disrupted Only in Male Survivors of CA/CPR

To assess whether amygdala function is altered in survivors of CA/CPR-induced GCI we incorporated the delay-fear conditioning learning and memory behavioral task (Figure 1A), which is well accepted as an amygdala dependent learning task (Naber & Bullinger, 2018; Yajie Sun et al., 2020). Both male and female animals that underwent the CA/CPR or sham procedure 7-days prior were taught to associate an innocuous CS (tone) with an aversive US (foot shock) in Context A (training). During training, baseline freezing was negligible in male and female animals and was not significantly different between sham and CA/CPR animals (Males: sham - n = 12, 6.181 ± 6.2% freezing; CA/CPR – n = 11, 0.121 ± 0.4% freezing; F _Time x Injury Status_ (1,21) = 5.53, p = 0.0.029; F _Time_ (1,21) = 205.8, p < 0.0001; F _Injury status_ (1,21) = 17.78, p = 0.33; F _mouse_ (21,21) = 1.208, p = 0.33 | Females: sham – n = 14, 3.899 ± 7.14% freezing; CA/CPR – n = 15, 6.700 ± 7.5% freezing; F _Time x Injury Status_ (1,27) = 11.61, p = 0.0021; F _Time_ (1,27) = 222.7, p < 0.0001; F _Injury Status_ (1,27) = 2.144, p = 0.15; F _mouse_ (27,27) = 2.369, p = 0.014). However, during the tone/shock pairing both male (sham, n = 12, 50.483 ± 12.9% freezing; CA/CPR, n = 11, 31.94 ± 12.2% freezing; t _sham v CA_ (1, 21) = 4.702, p < 0.0001, Fisher’s LSD) and female (sham, n = 14, 48.41 ± 14.43% freezing; CA/CPR, n = 15, 34.66 ± 16.07% freezing; t _sham v CA_ (54) = 3.084, p = 0.0032, Fisher’s LSD) CA/CPR survivors displayed diminished freezing, relative to sham animals (Figure 1B). 24-hours after training (day 2), mice were re-exposed to context A to evaluate hippocampal dependent spatial (contextual) memory. In males, there was a significant deficit in contextual fear memory in CA/CPR (n = 11, 47.23 ± 22.97% freezing; F _sex x injury status_ (1,48) = 1.192, p = 0.28; F _sex_ (1,48) = 1.811, p = 0.1847; F _injury status_ (1,48) = 13.03, p = 0.0007) animals when compared to shams (n = 12, 74.06 ± 9.2% freezing, q (48) = 4.451, p = 0.015, Tukey’s). However, in females, there was not a significant deficit in contextual fear memory (Sham, n = 14, 60.15 ± 22.71% freezing; CA/CPR, n = 15, 45.78 ± 22.51% freezing, q (48) = 2.678, p = 0.24, Tukey’s) (Figure 1C). 48-hours later (day 3) mice were placed in a novel context and freezing behavior was quantified during and after presentation of the CS as an indication of the animal’s ability to learn the CS-US association. Sham male animals (n = 12) displayed robust freezing to the CS in context B (52.44 ± 18.4% freezing; F _time x injury status_ (1,48) = 8.578, p = 0.008; F _time_ (1,21) = 79.97, p < 0.0001; F _injury status_ (1,21) = 8.657, p = 0.003; F _mouse_ (21,21) = 3.5, p = 0.003) as did sham female animals (n = 14; 42.1 ± 21.3% freezing; F _time x injury_ (1,27) = 0.69, p = 0.41; F _time_ (1,27) = 59.17, p < 0.0001; F _injury status_ (1,27) = 1.292, p = 0.27; F _mouse_ (27,27) = 1.65, p = 0.099) (Figure 1D). However, after CA/CPR, male animals were significantly diminished in their ability to form the CS-US association (n = 11; 26.71 ± 16.23% freezing; t (42) = 3.98, p = 0.003; Fisher’s LSD) when compared to sham males (Figure 1D). Interestingly, female survivors of CA/CPR were not diminished in their ability to form the CS-US association (n = 15; 33.22 ± 21.6% freezing; t (54) = 1.408, p = 0.165; Fisher’s LSD) (Figure 1D). The data shows male and female CA/CPR survivors display less freezing behavior during the acquisition of the CS/US association (training), however, only male animals are diminished in their ability to recall contextually aversive memories (context A testing) and in their ability to recall the CS/US association in a novel context (context B testing).

**Figure 1:**
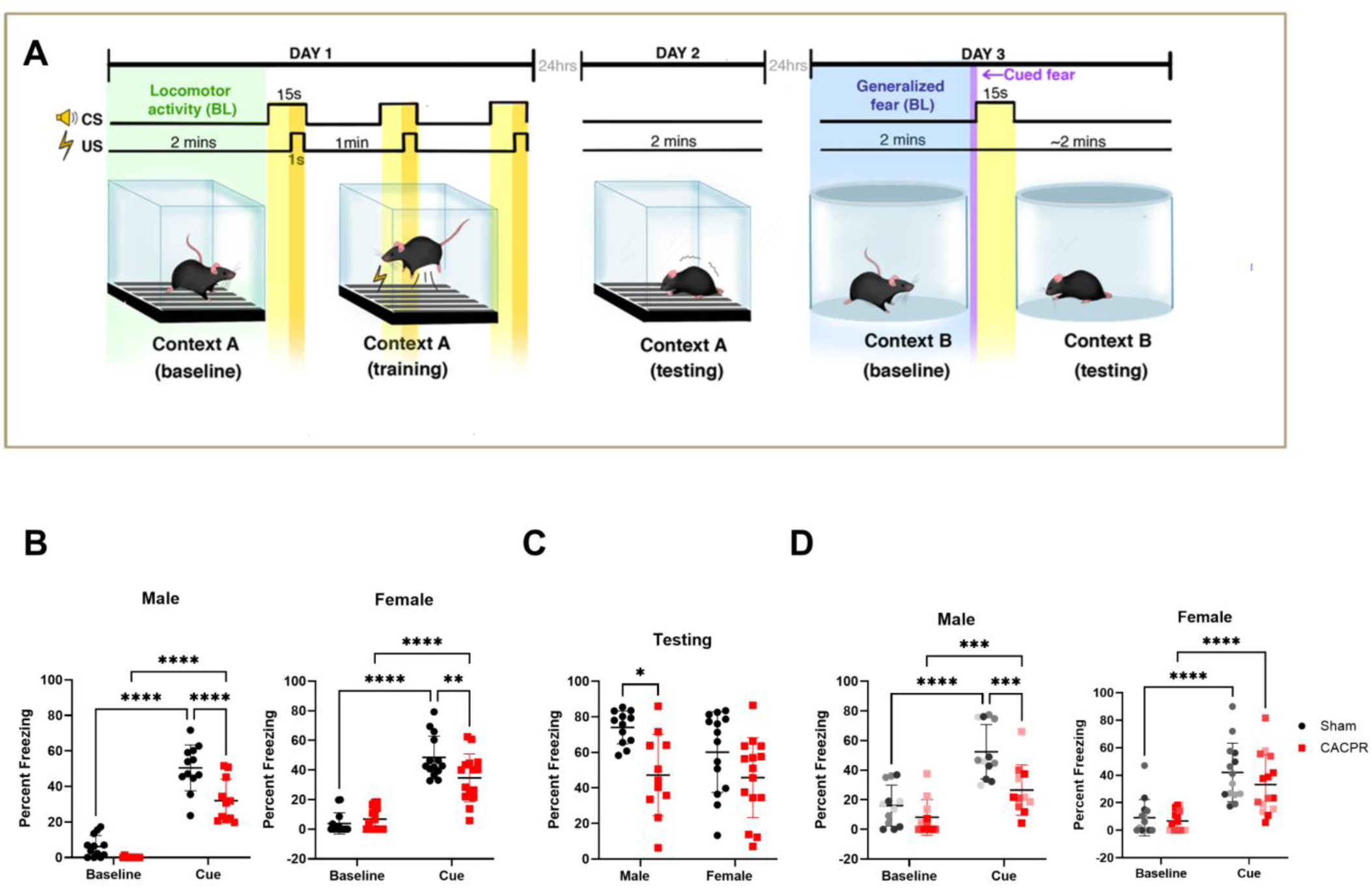
Only male survivors of CA/CPR are diminished in their ability to form associative memories. An illustration of the delay fear conditioning procedure performed in male and female mice (A). Quantification of the time spent freezing during training on Day 1 in male (left) and female (right) mice (B). Quantification of freezing behavior when the mice were re-introduced to context A on Day 2 (C). Quantification of the time spent freezing in response to presentation of the cue (CS) in a novel context in male (left) and female (right) mice (D). Two-way ANOVA (injury status x time point); * = p < 0.05, ** = p < 0.01, *** = p < 0.001, **** = p < 0.0001.

### LTP of the Cortical Input to the BLA but not the Intra-Amygdala Pathway is Disrupted in Male Mice That Survive CA/CPR

The requirement of LTP of the cortical input to the BLA (Supplemental Figure 1a) for the acquisition of associative memory is well documented(Luchkina & Bolshakov, 2019; Maren, 2017; Tsvetkov et al., 2002). Thus, it was necessary to record LTP in this pathway to gain further insight into GCI-induced amygdala dysfunction. We performed extracellular field potential recordings of the cortical input to the BLA in female and male mice 7-days after CA/CPR (Figure 2 A and E respectively). We found that female (n = 11, 136.0 ± 21.8% of BL) and male (n = 8, 143.6 ± 14.4% of BL) sham mice have robust potentiation of this pathway following theta-burst stimulation. In CA/CPR animals female LTP was not significantly different from sham females (n = 10, 130.4 ±23.9% of BL; t (19) = 0.56 (p = 0.58) but in male CA/CPR animals, LTP was significantly reduced (n = 10, 110.9 ± 14.98% of BL; t (16) = 4.68, p = 0.0003) (Figures 2A, B, E, and G). This technique also allows us to acquire information about other aspects of amygdala function such as excitability (input/output curve) measurements and estimates of pre-synaptic release probability (Paired Pulse Ratio-PPR). In female mice, we found that there is no significant GCI induced change in excitability, as indicated by comparison of the slope of the Input/Output curve (t (18) = 0.062, p = 0.95) (Figure 2C and Supplemental Figure 2b). We also found no GCI-induced change in excitability in male animals (t (11) = 0.898, p = 0.39) (Figure 2G and Supplemental Figure 1a). However, we did find a small but significant decrease in the paired pulse ratio (PPR) in female mice indicating a GCI induced increase in release probability (Sham: n = 12, PPR = 1.41 ± 0.21; CA/CPR: n = 10, PPR = 1.24 ± 0.14; t (20) = 2.196, p = 0.04) (Figure 2D). In male animals, we found that there was no significant difference in the PPR, as an indication of pre-synaptic release probability (Sham: n = 8, PPR = 1.38 ± 0.25; CA/CPR: n = 10, PPR = 1.32 ± 0.28; t (16) = 0.48, p = 0.63) (Figure 2H). Thus, our data suggest that there is male specific impairment of LTP that corresponds with the deficit in the ability to acquire associative memories. It also indicates that female CA/CPR survivors do not have impairment in amygdala dependent behavior or LTP, despite a GCI-induced change in PPR.

**Figure 2:**
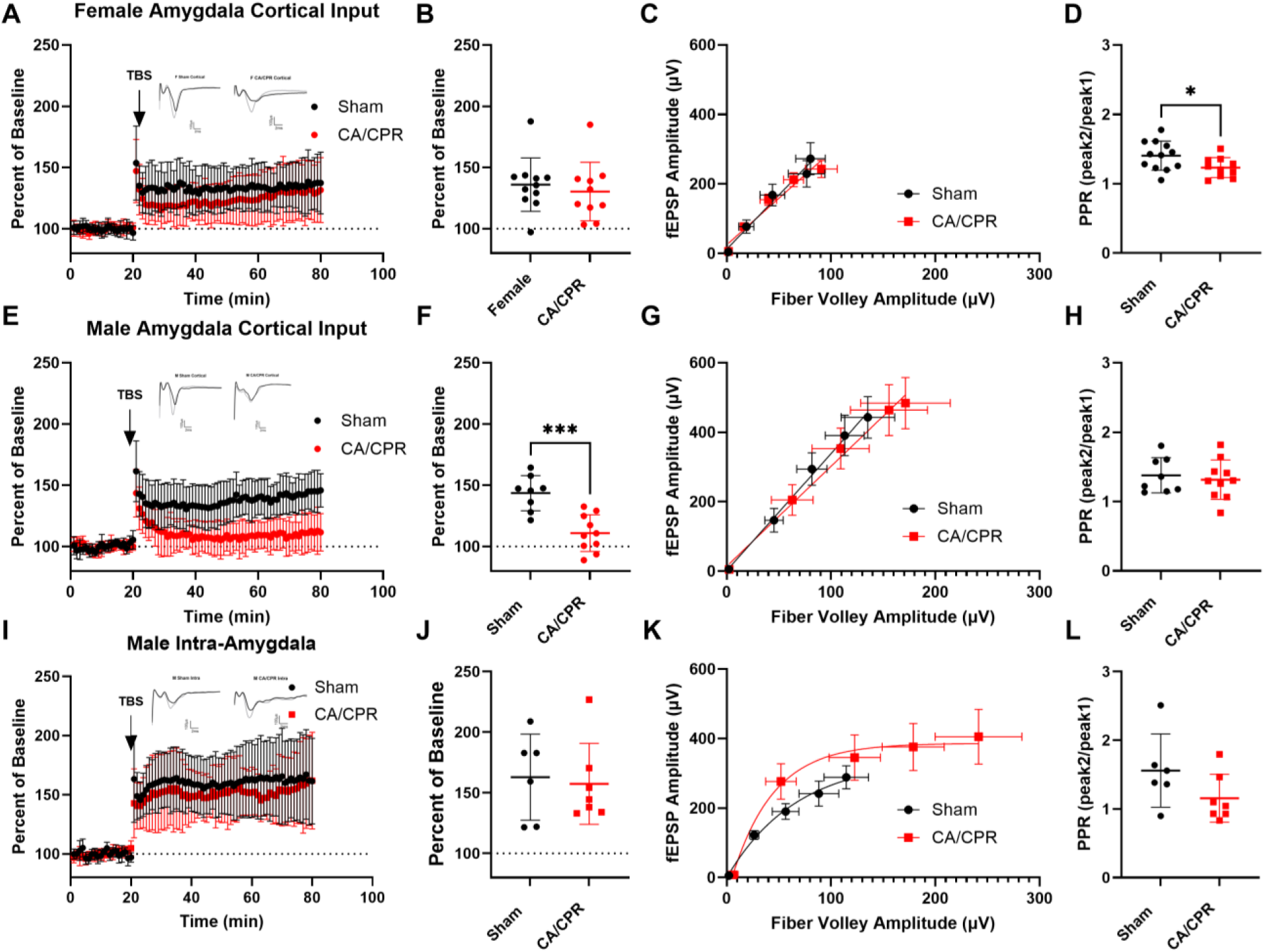
LTP in the cortical input to the BLA is disrupted by GCI while the intra-amygdala pathway is not. Time course of LTP recording of cortical input to the BLA in female mice (A). Quantification and comparison of last 10 minutes of LTP recordings (B) performed in A. Input/output curve for the recordings performed in A (C). Paired pulse ratio of data obtained in A (D). Time course of LTP recording of cortical input to BLA in male mice (E). Quantification and comparison of last 10 minutes of LTP recordings (F) performed in E. Input/output curve for the recordings performed in E (G). Paired pulse ratio of data obtained in E (H). Time course of Intra-amygdala LTP experiments (I). Quantification of last 10 minutes of LTP recordings (J) performed in I. Input/output curve for the recordings performed in I (K). Paired pulse ratio data obtained in I (L). Insets in A, E, and I display representative traces from sham (left) and CA/CPR (right) animals. Students T-test for B, D, F, H, J, and L. * = p < 0.05, *** = p < 0.001. C and G were fit with a simple linear regression. K was fit with a single exponential.

Given the male specific behavioral deficit and corresponding LTP deficit in the BLA, we further sought to assess LTP in a downstream amygdala pathway in male animals. This allows us to determine if this is a circuit specific effect or if the entire amygdala is dysfunctional in its ability to undergo LTP after GCI. The stimulating and recording electrodes were positioned as in Supplemental Figure 1b to record LTP of the intra-amygdala circuit. Interestingly, we found no significant difference in the magnitude of LTP in this circuit between sham (n = 6, 162.9 ± 35.5% of BL) and CA/CPR (n = 7, 157.4 ± 33.3% of BL, t (11) = 0.29, p = 0.78) (Figure 2I, J) animals. There was, however, a significant GCI-induced increase in excitability as evidenced by a significant increase in Input/Output curve (t (6) = 5.52, p = 0.0015) (Figure 2K and Supplemental Figure 2c). There was also no GCI-induced alteration in PPR (Sham: n = 6, PPR = 1.56 ± 0.53; CA/CPR: n = 7, PPR = 1.16 ± 0.35; t (11) = 1.64, p = 0.13) (Figure 2L). Despite the GCI-induced disruption of LTP in the cortical input to the BLA, LTP of the intra-amygdala circuit remains intact but more responsive to electrical stimuli. Given the male specific deficits in DFC and LTP after GCI we decided to only incorporate male animals into the rest of the study to further understand the mechanism of amygdala dysfunction rather than the sex difference in susceptibility to amygdala dysfunction after GCI.

### Associative Learning and Memory Deficit in Male Mice is Not Due to Locomotor Dysfunction

To assess the possibility of locomotor dysfunction in male survivors of CA/CPR, we incorporated the Open Field (OF) test in sham or CA/CPR male mice (Figure 3A, B). Additionally, the OF test can be used to assess another amygdala function, anxiety like behavior (Seibenhener & Wooten, 2015). Thus, in a separate cohort of animals, 7-days after the CA/CPR or sham procedure mice were placed into the OF apparatus and their behavior monitored and recorded for 10-minutes. The average speed and distance traveled by the animals was quantified in the OF apparatus. In male CA/CPR survivors, there was no difference in the distance traveled (sham 28.14 ± 7.984m vs. CA/CPR 25.92 ± 8.3m; t (28) = 0.73, p = 0.47) or average speed (sham 0.05806 ± 0.012 m/s vs. CA/CPR 0.061 ± 0.022; t (28) = 0.45, p = 0.6542) (Figure 3C, D). Thus, the GCI-induced associative learning and memory deficit reported above is not due to locomotor dysfunction.

**Figure 3:**
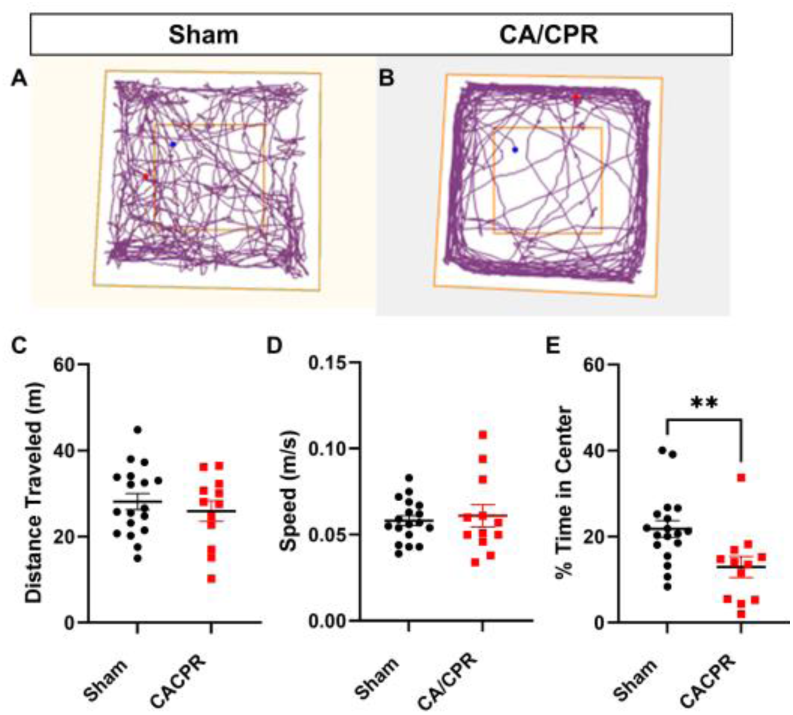
Locomotor dysfunction is not an underlying cause of the male specific DFC deficit. Representative path taken by a sham (A) animal and a CA/CPR (B) animal during the open field task. Quantification of the total distance traveled (C) and average speed (D) of each animal during the task. Percent time spent in the center of the apparatus, as an indication of anxiety like behavior

Rodents prefer to avoid open spaces and are deemed to be less anxious if more time is spent in the center of the OF apparatus. Male (n = 10) survivors of CA/CPR displayed anxiety like behavior when compared to shams of the same sex (male n = 21). Sham animals spent an average of 21.81 ± 8.2% of the time in the center of the apparatus and CA/CPR animals spent an average of 12.9 ± 8.5% of the time in the center of the apparatus (t (28) = 2.86, p = 0.0079) (Figure 3E). Given these results, it is not likely that locomotor dysfunction is contributing to either behavioral measure, the male specific DFC deficit, or the presence of anxiety like behavior. Interestingly, the result of increased anxiety like behavior in male CA/CPR survivors corresponds with the observation that male survivors also have increased excitability in the intra-amygdala circuit (Rau et al., 2015a; Rosen & Schulkin, 2022).

### Neurodegeneration in the Basolateral Amygdala is Not Contributing to the Observed Behavioral Dysfunctions in Male Mice

There is well documented neurodegeneration in select brain regions after GCI, including the hippocampus(Kofler et al., 2004; Quillinan et al., 2015; Rumian et al., 2021). However, throughout our literature search, we were unable to find any assessment of cell death in the amygdala after GCI. Thus, we used the FluoroJade (FJ) stain to quantify GCI-induced cell death in the amygdala, which could underlie the observed behavioral and LTP deficits. We found no FJ positive neurons in the amygdala at either 3 or 7 days after recovery from CA/CPR (Figure 4A, B, respectively). As an additional measure of cell death, we also quantified the number of NeuN^+^ cells in the BLA. There was no difference between sham (n = 3,118.7±18.72 cells) and CA/CPR (n = 4, 111.9±21.16 cells) animals (t (5) = 0.44, p = 0.6775) (Figure 4C). Thus, neurodegeneration within the amygdala is not contributing to the GCI induced behavioral deficits or the alterations in LTP or excitability. To confirm FJ staining was active, we assessed neuronal cell death in the hippocampus of the same animals, observing profound neurodegeneration in the hippocampus 3 days post CA/CPR that was diminished, but still present 7 days after CA/CPR (Supplemental Figure 3).

**Figure 4:**
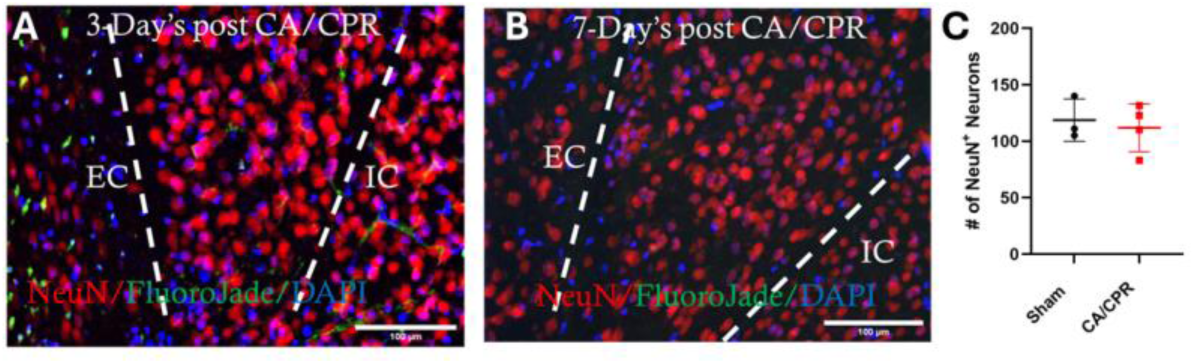
Neurodegeneration within the BLA is not contributing to the DFC deficit. Representative image of the BLA 3-days after CA/CPR (A). Representative image of BLA 7-days after CA/CPR (B). Quantification of the number of NeuN positive neurons within the BLA 3-days after CA/CPR (C). Students T-test.

### NMDA Receptors are not contributing to the circuit specific LTP deficit in male survivors of CA/CPR

A major contributor to the induction of LTP in many brain regions are the N-Methyl-D-Aspartate receptors (NMDAr) (Bin Ibrahim et al., 2022; Chen et al., 2021; Li et al., 2019), making reduction in NMDAr function a likely contributor to the impaired plasticity we observed in the amygdala. It has been demonstrated that during LTP recordings, the area of the fEPSPs during the TBS can be used to assess NMDAr function (Larson & Munkacsy, 2015). Thus, we were able to evaluate NMDAr function using our fEPSP recordings by quantifying the area of the fEPSP response during TBS stimulation (Figure 5A). EPSP area values were normalized to the area of the first burst in the TBS train and plotted in Figure 5B. There is no significant difference in the magnitude of the second burst relative to the first between sham (n = 8, 106.4 ± 9.6% of 1^st^ burst) and CA/CPR animals (n = 10, 108.2 ± 24.1% of 1^st^ burst; F _burst x injury status_ (1,16) = 0.039, p =0.85; F _burst_ (1,16) = 2.59, p = 0.13; F _injury status_ (1,16) = 0.039, p = 0.85; F _mouse_ (1,16) = 1.00, p = 0.5) fEPSP area during the induction of LTP (Figure 5C). Thus, it is not likely that dysfunctional NMDA receptors are contributing to the associative learning and memory deficit or the corresponding LTP deficit observed in the cortical input to the BLA.

**Figure 5:**
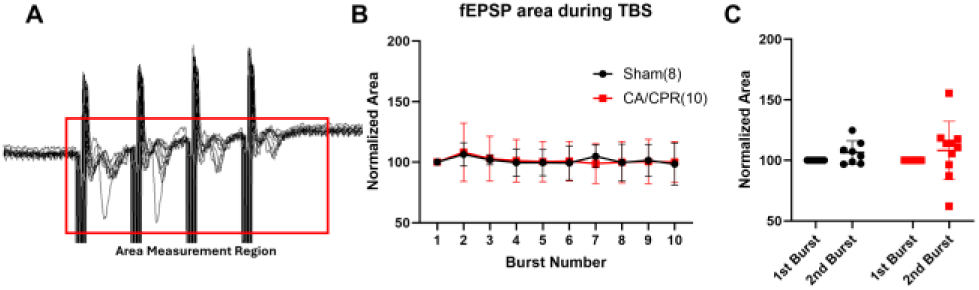
NMDA receptor function is not disrupted during TBS. Representative trace of the fEPSP response to TBS stimulation, red rectangle indicated the region used to acquire area measurements (A). Quantification of area during TBS stimulation normalized to the area of the first burst in the train (B). Boxplot of data from first and second burst in the during TBS (C). 2-Way ANOVA (injury status x burst number).

### L-Type Calcium Channel Function is minimally Altered in Male Survivors of CA/CPR

There is evidence of mechanistic differences in the induction of LTP between these two circuits we studied (cortical input to BLA or intra-amygdala). Drephal et al., 2006, and our own pharmacological experiments (Supplemental Figure 4) demonstrate the requirement of functional NMDA receptors in both circuits for the induction of LTP, however only the cortical input to the BLA requires functional L-type calcium channels (LTCCs) for induction of LTP.

Thus, we directly measured current responses mediated by LTCCs in pyramidal neurons in the BLA to evaluate the possibility that dysfunctional LTCCs are contributing to both the LTP deficit in the cortical input to the BLA and the deficit in DFC that we have observed. In visually identified BLA pyramidal neurons, whole-cell calcium currents were pharmacologically isolated (see material and methods) and normalized to cell capacitance to yield current density values that were used for comparison. The normalized calcium currents acquired during treatment with nifedipine (Figure 6B) were subtracted from the calcium currents acquired before perfusion of Nifedipine (Figure 6A). This subtracted value can therefore be attributed to LTCC mediated calcium current (Figure 6C). There was a significant difference in the total calcium current between sham and CA/CPR neurons at select holding potentials (F _current x injury status_ (9,135) = 1.88, p = 0.061; F _current_ (1.375, 20.62) = 46.6, p < 0.0001; F _injury status_ (1,15) = 2.4, p = 0.14; F _mouse_ (15, 135) = 10.27, p < 0.0001; Tukey’s post hoc: −50mV, q(12.18) = 3.6, p = 0.03; −40mV, q(11.8) = 4.49, p = 0.008; +20mV, q(14.98) = 3.04, p = 0.049; +30mV, q(14.18) = 4.29, p = 0.009; and +40mV, q(12.45) = 5.09, p = 0.01) (Figure 6A). There was also a significant difference in the nifedipine sensitive current at two holding potentials (F _current x injury status_ (9, 144) = 1.75, p = 0.08; F _current_ (1.075,17.2) = 2.159, p = 0.16; F _injury status_ (1,16) = 3.93, p = 0.065; F _mouse_ (16,144) = 2.17, p = 0.008; Tukey’s post hoc: −30mV, q (10.07) = 3.37, p = 0.038; and +20mV, q (9.33) = 3.22, p = 0.048) (Figure 6B). However, when the subtraction was performed, there was only a significant difference between sham and CA/CPR animals at a holding potential of +20mV (F _current x injury status_ (9,144) = 0.88, p = 0.55; F _current_ (2.009,32.14) = 25.52, p < 0.0001; F _injury status_ (1,16) = 0.92, p = 0.35; F _mouse_ (16,144) = 2.67, p = 0.001; Tukey’s post hoc: Sham: n = 9, −10.17 ± 6.54pA/pF, CA/CPR: n = 9, −2.5 ±2.79pA/pF; q (10.8) = 4.57, p = 0.008). This effect, at such a highly depolarized potential is likely minimally influencing cellular function considering the cell likely only reaches this voltage during action potentials. However, the observed differences do suggest a GCI-induced alteration in calcium signaling in BLA pyramidal neurons. Our results also suggest that dysfunction of LTCCs has little if any role in this altered calcium signaling. It is likely that GCI-induced alterations of calcium dynamics in BLA pyramidal neurons is contributing to the observed deficit of LTP in the cortical input to the BLA and is therefore disrupting associative learning and memory processes that depend on the amygdala.

**Figure 6:**
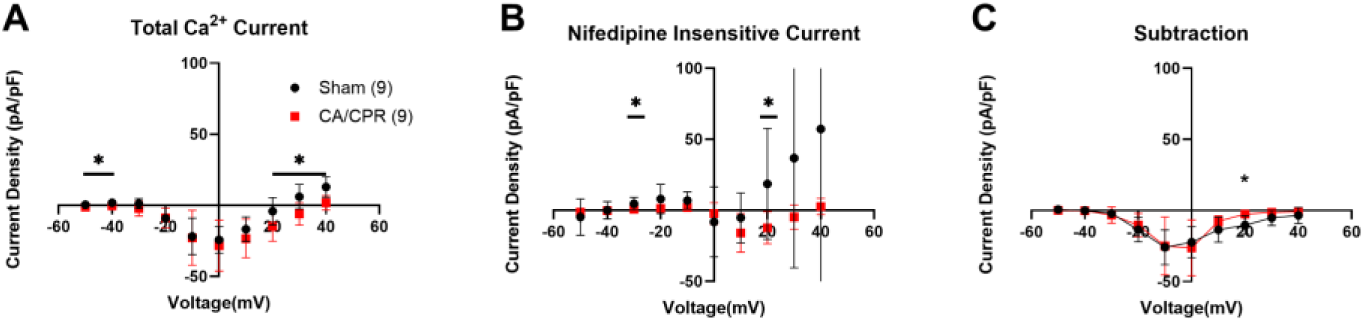
L-type calcium channels and NMDA receptors are not contributing to the LTP deficit. Pharmacologically isolated whole-cell calcium currents in response to voltage steps in BLA pyramidal neurons (A). Whole cell current response of the currents recorded in A after perfusion of the LTCC blocker Nifedipine (B). Current attributed to LTCCs in BLA pyramidal neurons after subtracting the current values in B from those in A. Two-way ANOVA (Injury Status x Current), * = p < 0.05.

### Calcium Signaling is Increased in Distal Dendritic Spines of Pyramidal Neurons in the Basolateral Amygdala

To further evaluate the possibility that calcium signaling is disrupted in BLA pyramidal neurons after GCI we combined whole-cell electrophysiology with 2-Photon calcium imaging. Calcium indicator (Cal520) and cell fill (Alexa 594) were loaded into BLA pyramidal neurons via the patch pipette and a glass, monopolar, stimulating electrode was placed near the secondary dendritic branch of interest. A current pulse was delivered to elicit a voltage response in the patched cell to ensure synaptic communication. A representative image of the positioning of the recording and stimulating electrode is presented in Figure 7A and a higher magnification image representing the positioning of the stimulating electrode relative to the branch of interest is presented in Figure 7B. Line scans of the calcium response to electrical stimulation and back-propagating action potentials (bAP) were then acquired in individual dendritic spines for comparison (Figure 7C, D).

**Figure 7:**
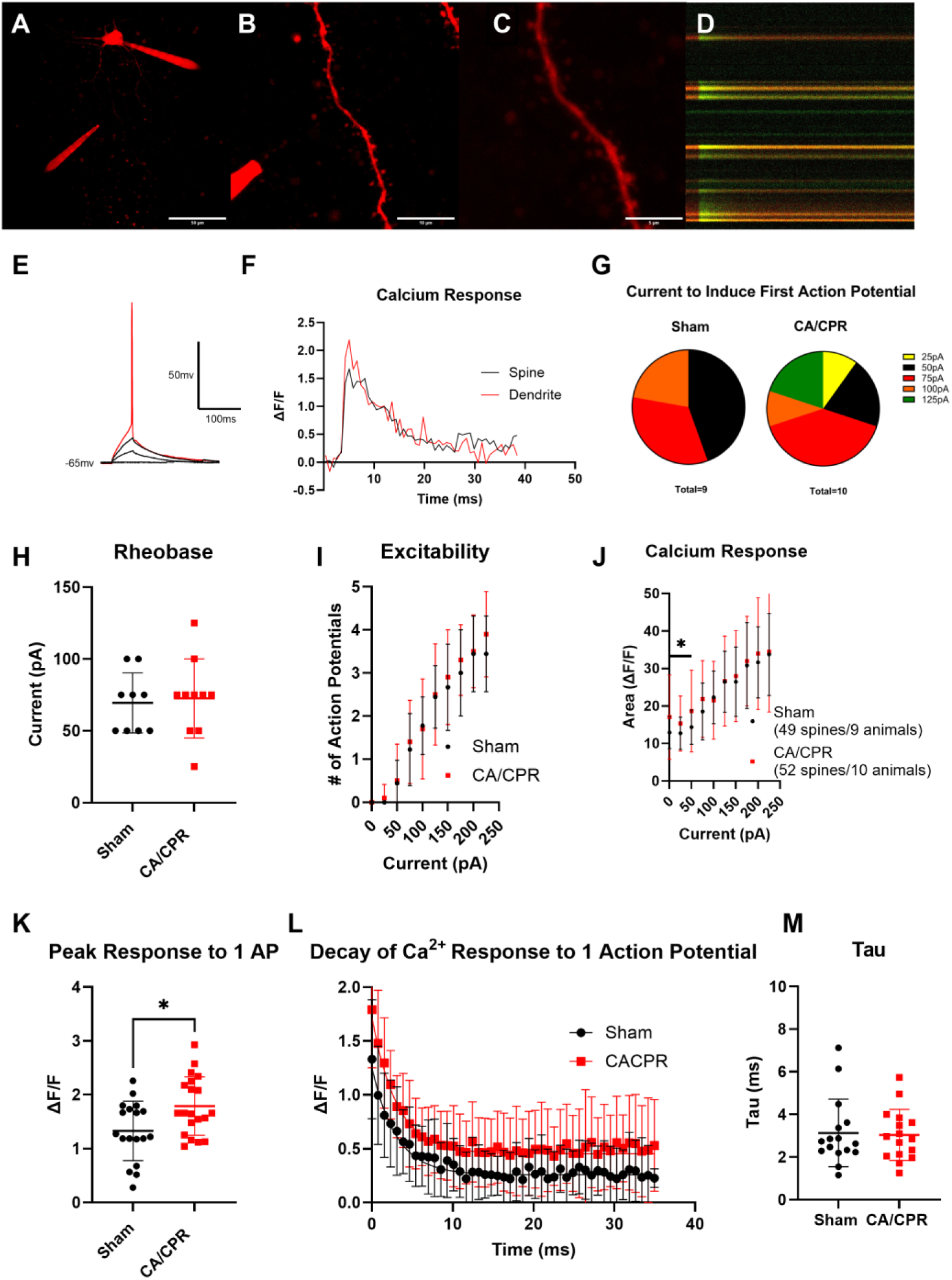
Experimental design and results to evaluate the contribution of distal dendritic spine calcium signaling to LTP and DFC deficits in GCI survivors. Representative Z-projection of entire neuron including location of stimulation electrode relative to the soma and the patch pipette that is delivering the cell fill and calcium indicator (A). 3x zoom of the same neuron in A, showing the spines and dendrite of interest and the position of the stimulating electrode relative to the spines (B). Representative image depicting the location of line scans used to image calcium transients during intrinsic depolarizations and synaptic transmission (C). Representative image of line scans acquired during a single back propagating action potential, green is calcium response and red is cell fill (D). Single action potential that resulted from delivery of a 75pA depolarizing current pulse (E) and the resulting calcium transiens produced in the dendrite and spine of interest (F). Pie charts depicting the amount of depolarizing current required to induce a single action potential in BLA pyramidal neurons (G). Comparison of the whole-cell properties of rheobase (H) and excitability (I). Plot of the area under the curve of the calcium responses induced by depolarizing current steps (J). Plot of the peak of the calcium response in subset of neurons that only produced one action potential at rheobase (K). Visualization of the decay phase of the calcium responses in K (L). Comparison of the Tau constants that correspond to the decay phase of the responses in L (M). Mann-Whitney test for H, Nested ANOVA for K and M, Two-way ANOVA (current x action potential number or area) for I and J. * = p < 0.001.

We quantified the calcium response in individual distal dendritic spines to bAPs, which is reported to be mediated by voltage-gated calcium channels (Theis et al., 2018). Brief current steps were delivered to the patched neuron and the voltage response was recorded (Figure 7E), to evaluate intrinsic excitability of the neurons, concurrently with the corresponding calcium response in individual dendritic spines (Figure 7F), expressed as ΔF/F for comparison between groups. There was some variability in the number of action potentials produced at rheobase. Figure 7G shows the current at which rheobase occurs and shows that in many neurons this is between 25-75pA. We found that there is no significant difference in mean rheobase (Sham: 69.44 ± 20.8pA; CA/CPR: 72.5 ± 27.5pA, U = 41.5, p = 0.82) (Figure 7H), or excitability (F _current x injury status_ (9,153) = 0.43, p = 0.92; F _current_ (1.48,25.23) = 143, p < 0.0001; F _injury status_ (1,17) = 0.2, p = 0.66; F _cell_ (17,153) = 15.67, p < 0.0001; Sigmoidal fit Hill slope: Sham 2.403, CA/CPR 2.067; F(4, 182) = 0.54, p = 0.71) between sham or CA/CPR BLA pyramidal neurons (Figure 7I). However, there was significantly higher basal calcium in CA/CPR animals when 0pA-50pA of current were delivered (F _current x injury status_ (9,891) = 1.64, p = 0.1; F _current_ (2.147,212.6) = 130.5, p < 0.0001; F _injury status_ (1,99) = 1.43, p = 0.24; F _spine_ (99,891) = 15.22, p < 0.0001; Tukey’s post hoc: 0pA-Sham: 12.96 ± 4.3, CA/CPR: 17.03 ± 11.3, q (66.2) = 3.44, p = 0.018; 25pA-Sham: 12.72 ± 4.3, CA/CPR: 15.31 ± 7.3, q(83.41) = 3.08, p = 0.032; 50pA-Sham: 14.37 ± 4.6, CA/CPR: 18.67 ± 10.9, q(69.36) = 3.68, p = 0.011) to the soma of the neuron through the patch pipette (Figure 7J). To ensure consistency in our comparison of calcium transients to bAP, we only included cells that produced one action potential at rheobase (n = 9 Sham, 10 CA/CPR) for spine calcium transient comparisons. After selection of cells that produced one action potential at rheobase, we were able to quantify and compare peak calcium response in 18 spines from sham animals and in 20 spines from CA/CPR animals. There was a significant increase in calcium response to one bAP in BLA pyramidal neurons after GCI produced by CA/CPR (ΔF/F: Sham: 0.33 ± 0.55, CA/CPR: 1.79 ± 0.54, F _injury status_ (1,14) = 11.179, p = 0.0031; F _spine_ (14,21) = 3.715, p = 0.0034) (Figure 7K). It is possible that calcium buffering in the spines is disrupted by GCI. So, we decided to also look at the decay kinetics of the calcium response. Overall, BLA neurons in CA/CPR animals had elevated calcium response for the entire time course of the recordings, however the shape of the decay response was similar between groups (Figure 7L) as was the Tau constant when the decay responses were fit with a single-phase exponential decay function (Sham: 3.13 ± 1.6ms, CA/CPR: 3.04 ± 1.2ms; F _injury status_ (1,13) = 0.13, p = 0.72; F _spine_ (13,17) = 8.32, p < 0.0001) (Figure 7M). This result shows that after GCI, BLA pyramidal neurons are not altered in their intrinsic electrophysiological properties, however, the intrinsic calcium response to back propagating action potentials is significantly increased after GCI.

In addition to the intrinsic calcium signaling properties of BLA pyramidal neurons after GCI, we also sought to identify any dysfunction in synaptic calcium signaling. To this end, we delivered TBS, as in the previously performed LTP experiments (Figure 2) and recorded the calcium response to electrical LTP induction stimuli. When TBS was delivered (Figure 8A), many spines displayed a spiking pattern of calcium transients that corresponded to the burst of electrical stimuli that were delivered (Figure 8B). The calcium responses were quantified and expressed as ΔF/F (Figure 8C). The area of the calcium response in each spine was then calculated in Prism (Graph Pad, San Diego, CA). These calculated area values were then statistically compared. We found that in CA/CPR survivors, the distal dendritic spines of BLA pyramidal neurons have a larger calcium response to electrical induction of LTP (n = 46 spines from 9 animals; 102.2 ± 53.6 (ΔF/F)/ms,) than the sham control group (n = 54 spines from 9 animals; 81.68 ± 34.53 (ΔF/F)/ms; F _injury status_ (1,16) = 8.64, p = 0.004; F _spine_ (16/77) = 5.68, p < 0.0001) (Figure 8D). Thus, not only is intrinsic calcium signaling elevated in BLA pyramidal neurons after GCI, but calcium signaling during electrical induction of LTP is also elevated. This GCI-induced alteration of calcium dynamics is likely underlying the observed LTP deficit and corresponding dysfunction of associative learning and memory processes.

**Figure 8:**
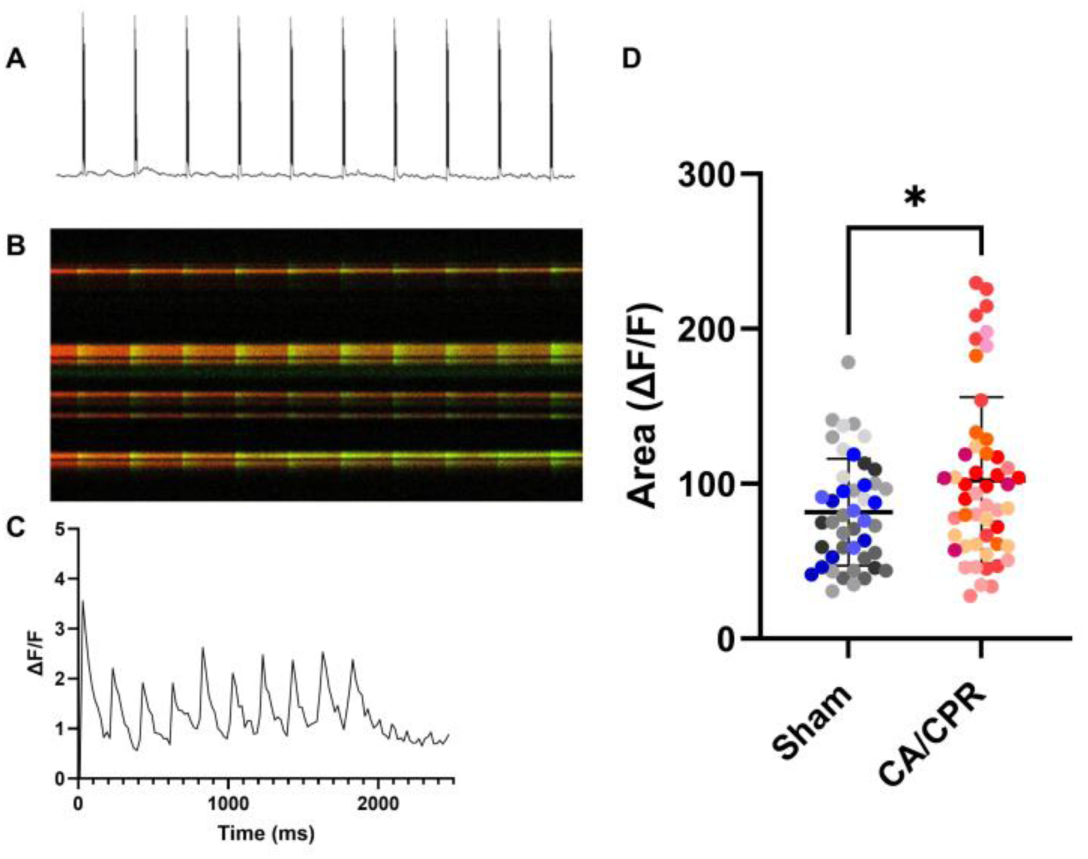
Increased calcium signaling after GCI is also observed during the synaptic induction of LTP. An LTP inducing stimuli (TBS) was delivered through the stimulating electrode (A), which produced waves of calcium transients that correspond with the electrical stimuli (B) and is quantified by calculating ΔF/F (C). Plot depicting the area under the curve of the response of the calcium transients to electrical stimuli (D). Data points of the same shade of color are individual spine measurements from the same animal. Nested ANOVA, * = p < 0.05.

## Discussion

We found that in survivors of CA/CPR-induced GCI, there is male-specific impairment of amygdala function that persists for at least 7-days after injury. Discovery of this amygdala dysfunction prompted us to pursue a greater understanding of the mechanism of this male specific amygdala-dependent learning and memory deficit. Using LTP recordings we identified a corresponding male-specific deficit in a circuit that is essential for the acquisition of associative memories (Maren, 2005). Some likely mechanisms for this associative learning dysfunction in male mice, such as locomotor impairment or loss of neurons in the amygdala were excluded as possibilities. We found no evidence of neurodegeneration in the BLA and found no evidence of locomotor dysfunction in the OF task. We did however find that male CA/CPR survivors also display anxiety-like behavior and have increased synaptic excitability within the amygdala, a well validated feature of anxiety disorders (Rau et al., 2015b; Rosen & Schulkin, 2022). It appears that there is a GCI-induce deficit in the acquisition of associative fear learning, as there is reduced freezing during the CS/US pairing in the training phase of the experiment (day 1). However, this deficit was also present in female CA/CPR survivors without a corresponding deficit in contextual fear memory or cued fear memory (amygdala dependent). It is possible that female mice display a behavior other than freezing in response to foot-shock, as darting behavior in response to foot-shock has been reported in female rats during aversive fear learning (Mitchell et al., 2022). However, the lack of a deficit in freezing behavior in female CA/CPR survivors during testing in context A (day 2) is surprising, as our lab has published results showing similar deficits in contextual fear memory in male and female mice (Dietz et al., 2020) after GCI. It is important to note, however, that the behavioral paradigm used in this study incorporates a tone during training and our previous study did not. Thus, this current study utilized background contextual fear conditioning, and our previous studies utilized foreground contextual fear conditioning (shock without tone). There are mechanistic differences between the two paradigms(Phillips & Ledoux, 1994), and it is possible that GCI differentially affects the two types of learning in female mice, yielding deficits in foreground but not background contextual fear learning. Despite these anomalies in female mice, our premise of male specific amygdala dysfunction after GCI still stands given our results in the LTP experiments and the outcome of the freezing behavior in context B (day 3). Additionally, despite female mice not displaying any amygdala dependent behavioral or LTP deficits as a result of GCI, there was still a GCI induced change in pre-synaptic plasticity (PPR) in the BLA that was not further explored in this study.

There is clinical evidence to suggest that the outcome of CA/CPR is more favorable in females than similarly aged males(Fan, 2017; Fels & Manfredi, 2019; Siegel et al., 2010), at least until the onset of menopause, in terms of survival. However, due to lack of continued follow up of CA/CPR survivors we know very little about the effect that biological sex has on neurological outcome in this population. It is possible that the differential production of sex hormones between male and female survivors is influencing neurological outcome, and it is possible that this is also influencing the sex difference we observed in this study. However, we decided not to pursue the mechanism underlying the difference in susceptibility to amygdala dysfunction after GCI between male and female mice. Rather we directed our efforts to further understanding the mechanism of the observed male specific amygdala dysfunction after GCI. Our results further highlight the importance of incorporating biological sex as a variable in scientific studies and warrants future studies to determine the mechanism of female robustness to amygdala dysfunction after GCI, and if there might be other female-specific behavioral deficits that manifest because of the GCI-induced change in pre-synaptic release probability in the cortical input to the BLA that was revealed by our study.

We were unable to implicate NMDA receptors as the underlying cause of BLA dysfunction in males. Considering the role that NMDA receptors play in plasticity in both the cortical and intra-amygdala circuits and the presence of LTP dysfunction in only one of those circuits after GCI, it is reasonable that NMDA receptors play no role in plasticity impairments. However, we were not able to exclude the possibility that LTCCs could be playing a role in amygdala dysfunction in males. We did observe a small but significant change in LTCC current density after GCI, however, a larger effect could be masked by caveat of the technique, specifically, space clamp issues (Bar-Yehuda & Korngreen, 2008). There is incomplete voltage control over the entirety of the neuron due to the extensive branching in dendritic arbors and the axon. However, our results do suggest that calcium is playing a role in amygdala dysfunction after GCI in male animals.

Calcium is tightly regulated in neurons and any change in calcium dynamics will alter synaptic and intrinsic functions. The magnitude, duration, and localization of calcium responses all influence plasticity and are known to mediate directionality of plasticity, either LTP or LTD, and therefore learning and memory (Bauer et al., 2002; Chindemi et al., 2022; Evans & Blackwell, 2015; Nanou & Catterall, 2018; Yang et al., 1999). To more specifically evaluate synaptic alterations in calcium signaling after GCI we performed whole-cell electrophysiology experiments coupled with 2-photon calcium imaging in individual dendritic spines. We found that there is a significant increase of the calcium response in dendritic spines of BLA pyramidal neurons to a single bAP. The voltage changes produced in dendritic spines by bAP are necessary for the induction of LTP and are mediated by voltage gated calcium channels (VGCCs)(Cavazzini et al., 2005; Fuenzalida et al., 2010; Jaafari & Canepari, 2016). The composition of VGCCs that are mediating the calcium response to bAPs is not fully understood, but our result leaves open the possibility that LTCCs are still playing a role in the male-specific BLA dysfunction we have discovered. It also suggests the possibility that other VGCCs could be playing a role as well, such as T-type and R-type calcium channels, as both have been implicated in propagating the calcium response during bAPs (Kampa et al., 2006; Yuste et al., 1999). Future studies should incorporate pharmacological agents to determine which VGCCs are altered by GCI and how restoring the bAP mediated calcium response to sham levels influences plasticity.

Given the synaptic nature of the LTP deficit that we discovered, we also assessed the calcium response to synaptic induction of LTP in individual dendritic spines in BLA pyramidal neurons. Using this method, we found that there is also increased calcium response to synaptic induction of LTP in male survivors of CA/CPR. The result appears contrary to accepted mechanistic understandings of LTP induction, which require elevated calcium levels to activate intracellular signaling cascades that are responsible for induction and maintenance of LTP (Malenka, 1991; Simons et al., 2009). However, calcium dysregulation resulting in elevated calcium responses has been implicated in early stages of other learning and memory dysfunctions, such as Alzheimer’s Disease(McDaid et al., 2020). Additionally, aged neurons are known to have elevated calcium influx and reduced LTP that result from changes in VGCC expression(Oh et al., 2013). Thus, our observation of elevated calcium signaling in BLA pyramidal neurons is likely the cause of the LTP dysfunction and ultimately the behavioral deficits observed after GCI, as elevated calcium signaling does have the ability to disrupt LTP and learning and memory. While we were unable to directly implicate any specific source of calcium to these GCI-induced dysfunctions, we can narrow the possibilities based on our results and provide targeted directions for future research.

There are many calcium sources that contribute to synaptic and intrinsic cellular functions. It is not likely that calcium buffering or extrusion mechanisms are contributing to the dysfunction, as we found no change in decay kinetics of the calcium response to a bAP (Sabatini et al., 2002). We are unable evaluate the decay kinetics of the synaptic responses, but the mechanisms of extrusion and buffering of calcium from dendritic spines likely do not differ based on the source of depolarization (synaptic or intrinsic)(Franks & Sejnowski, 2002). We suggest two likely remaining possibilities: The first being changes in ion channel expression or function. We are not able to exclude the possibility that VGCCs are playing a role in synaptic dysfunction induced by GCI, and our results in the intrinsic calcium dynamics study suggest they are playing a role. It is also possible that the ischemic insult and resulting elevations of extracellular glutamate throughout the brain is causing ischemic LTP and insertion of calcium permeable AMPA receptors to the synapse (Kwak & Weiss, 2006; Noh et al., 2005; Orfila et al., 2018), however ischemic LTP has not been evaluated in the amygdala. This could explain our synaptic calcium increases, but not the intrinsic calcium increase, as there is low likelihood of AMPA receptor activation to bAP. If such a mechanism were discovered, it would indicate that the synaptic and intrinsic calcium signaling changes induced by GCI differ in mechanism. Another possibility is that there are GCI-induced changes in transient receptor potential (TRP) ion channels. TRP channels conduct calcium ions and have been implicated in neurodegeneration and changes in LTP after GCI, such as TRPM7, which has been implicated as a mediator of ischemic cell death, or TRPM2, which has a role in ischemic cell death and plasticity impairments after GCI (Dietz et al., 2020; Dietz et al., 2021; Hong et al., 2023; Turlova et al., 2021). The other possibility that could account for elevated dendritic spine calcium are changes in intracellular calcium store release. Calcium release from intracellular stores, such as mitochondria or endoplasmic reticulum, performs a vital role in intrinsic excitability and synaptic plasticity (Koek et al., 2024; Sandler & Barbara, 1999; Tedoldi et al., 2020; Valdés-Undurraga et al., 2023). Calcium-induced calcium release (CICR) from intracellular stores can be activated by either VGCCs or ligand activated ion channels and thus both synaptic and intrinsic calcium events can be modified by changes in CICR. CICR amplifies calcium responses in neurons and increased efflux of calcium from intracellular stores into the cytoplasm during depolarization (either synaptic or intrinsic) could account for our observations of increased calcium signaling. Both possibilities warrant further study to fully elucidate the underlying mechanism of this GCI induced increase in calcium signaling in BLA pyramidal neurons.

Overall, we implicate changes in calcium signaling in pyramidal neurons as the underlying mechanism of synaptic dysfunction in the BLA of male survivors of CA/CPR. The GCI-induced increase in calcium signaling, both intrinsically and synaptically, is disrupting LTP of the cortical input to the BLA and the ability for forming associative memories. Associative learning and memory processes are mediated by the amygdala and dysfunction of associative learning and memory has been implicated as the underlying cause of many affective disorders, including PTSD, anxiety, and depression(Allen et al., 2019; Diener, 2013; Grillon, 2002; Pittig et al., 2018). It is also worth noting that there are reported sex differences in the prevalence of neuropsychiatric disorders in humans, with PTSD, anxiety, and depression all being more prevalent in females (Peterlin et al., 2011; Seedat et al., 2009; Wickens et al., 2018). However, our results suggest that males are more susceptible to emotional dysfunction after ischemia. Thus, our study fills an important gap in knowledge about the functional outcome of GCI on the amygdala and highlights the need for extended follow up of human GCI survivors to accurately diagnose and treat affective disorders that may develop. We are the first to extensively study amygdala function after CA/CPR induced GCI and show that amygdala dysfunction after GCI is sex and circuit specific. Our study highlights the importance of sex as a biological variable and illustrates important differences in outcomes that are independent of neuronal cell death and provides mechanistic insights into how plasticity dysfunctions manifest differently based on sex.

## Acknowledgements

This work was supported by a Diversity Supplement (NS046072-S1) (JV), an AHA predoctoral fellowship award (23PRE1020271) (JV), an NIH Blueprint Diversity Specialized Predoctoral to Postdoctoral Advancement in Neuroscience (D-SPAN) award (F99NS135766-01) (JV), and RO1NS046072 (NQ/PSH).

## References

Allen, M. T., Myers, C. E., Beck, K. D., Pang, K. C. H., & Servatius, R. J. (2019). Inhibited Personality Temperaments Translated Through Enhanced Avoidance and Associative Learning Increase Vulnerability for PTSD [Review]. Frontiers in Psychology, 10. 10.3389/fpsyg.2019.00496

Anderson–Darling Test. (2008). In The Concise Encyclopedia of Statistics (pp. 12–14). Springer New York. 10.1007/978-0-387-32833-1_11

Anglada-Figueroa, D., & Quirk, G. J. (2005). Lesions of the basal amygdala block expression of conditioned fear but not extinction. J Neurosci, 25(42), 9680–9685. 10.1523/JNEUROSCI.2600-05.2005

Bar-Yehuda, D., & Korngreen, A. (2008). Space-clamp problems when voltage clamping neurons expressing voltage-gated conductances. Journal of Neurophysiology, 99(3), 1127–1136. 10.1152/jn.01232.2007

Bauer, E. P., Schafe, G. E., & Ledoux, J. E. (2002). NMDA Receptors and L-Type Voltage-Gated Calcium Channels Contribute to Long-Term Potentiation and Different Components of Fear Memory Formation in the Lateral Amygdala. The Journal of Neuroscience, 22(12), 5239–5249. 10.1523/jneurosci.22-12-05239.2002

Bin Ibrahim, M. Z., Benoy, A., & Sajikumar, S. (2022). Long-term plasticity in the hippocampus: maintaining within and ‘tagging’ between synapses. The FEBS Journal, 289(8), 2176–2201. 10.1111/febs.16065

Cavazzini, M., Bliss, T., & Emptage, N. (2005). Ca2+ and synaptic plasticity. Cell Calcium, 38(3-4), 355–367. 10.1016/j.ceca.2005.06.013

Chen, Q.-Y., Li, X.-H., & Zhuo, M. (2021). NMDA receptors and synaptic plasticity in the anterior cingulate cortex. Neuropharmacology, 197, 108749. 10.1016/j.neuropharm.2021.108749

Chew, H. S. J., Cheng, H. Y., Cao, X., & Lopez, V. (2018). Cognitive Assessment in Adult Cardiac Arrest Survivors: What Is Known and How Shall We Move Forward? Connect: The World of Critical Care Nursing, 12(3), 56–69. 10.1891/1748-6254.12.3.56

Chindemi, G., Abdellah, M., Amsalem, O., Benavides-Piccione, R., Delattre, V., Doron, M., Ecker, A., Jaquier, A. T., King, J., Kumbhar, P., Monney, C., Perin, R., Rössert, C., Tuncel, A. M., Van Geit, W., Defelipe, J., Graupner, M., Segev, I., Markram, H., & Muller, E. B. (2022). A calcium-based plasticity model for predicting long-term potentiation and depression in the neocortex. Nature Communications, 13(1). 10.1038/s41467-022-30214-w

Cohan, C. H., Neumann, J. T., Dave, K. R., Alekseyenko, A., Binkert, M., Stransky, K., Lin, H. W., Barnes, C. A., Wright, C. B., & Perez-Pinzon, M. A. (2015). Effect of Cardiac Arrest on Cognitive Impairment and Hippocampal Plasticity in Middle-Aged Rats. PLoS One, 10(5), e0124918. 10.1371/journal.pone.0124918

Diener, C. (2013). Altered Associative Learning and Learned Helplessness in Major Depression. In. InTech. 10.5772/51787

Dietz, R. M., Cruz-Torres, I., Orfila, J. E., Patsos, O. P., Shimizu, K., Chalmers, N., Deng, G., Tiemeier, E., Quillinan, N., & Herson, P. S. (2020). Reversal of Global Ischemia-Induced Cognitive Dysfunction by Delayed Inhibition of TRPM2 Ion Channels. Translational Stroke Research, 11(2), 254–266. 10.1007/s12975-019-00712-z

Dietz, R. M., Orfila, J. E., Chalmers, N., Minjarez, C., Vigil, J., Deng, G., Quillinan, N., & Herson, P. S. (2021). Functional Restoration following Global Cerebral Ischemia in Juvenile Mice following Inhibition of Transient Receptor Potential M2 (TRPM2) Ion Channels. Neural Plasticity, 2021, 1–10. 10.1155/2021/8774663

Drephal, C., Schubert, M., & Albrecht, D. (2006). Input-specific long-term potentiation in the rat lateral amygdala of horizontal slices. Neurobiol Learn Mem, 85(3), 272–282. 10.1016/j.nlm.2005.11.004

Elmer, J., & Callaway, C. W. (2017). The Brain after Cardiac Arrest. Semin Neurol, 37(1), 19–24. 10.1055/s-0036-1597833

Evans, R. C., & Blackwell, K. T. (2015). Calcium: Amplitude, Duration, or Location? The Biological Bulletin, 228(1), 75–83. 10.1086/bblv228n1p75

Fan, F. (2017). Menopause and Ischemic Stroke: A Brief Review. MOJ Toxicology, 3(4). 10.15406/mojt.2017.03.00059

Fels, J. A., & Manfredi, G. (2019). Sex Differences in Ischemia/Reperfusion Injury: The Role of Mitochondrial Permeability Transition. Neurochemical Research, 44(10), 2336–2345. 10.1007/s11064-019-02769-6

Franks, K. M., & Sejnowski, T. J. (2002). Complexity of calcium signaling in synaptic spines. BioEssays, 24(12), 1130–1144. 10.1002/bies.10193

Fuenzalida, M., Fernandez de Sevilla, D., Couve, A., & Buno, W. (2010). Role of AMPA and NMDA receptors and back-propagating action potentials in spike timing-dependent plasticity. J Neurophysiol, 103(1), 47–54. 10.1152/jn.00416.2009

Grillon, C. (2002). Associative learning deficits increase symptoms of anxiety in humans. Biol Psychiatry, 51(11), 851–858. 10.1016/s0006-3223(01)01370-1

Hong, D. K., Kho, A. R., Lee, S. H., Kang, B. S., Park, M. K., Choi, B. Y., & Suh, S. W. (2023). Pathophysiological Roles of Transient Receptor Potential (Trp) Channels and Zinc Toxicity in Brain Disease. International Journal of Molecular Sciences, 24(7), 6665. 10.3390/ijms24076665

Jaafari, N., & Canepari, M. (2016). Functional coupling of diverse voltage-gated Ca(2+) channels underlies high fidelity of fast dendritic Ca(2+) signals during burst firing. J Physiol, 594(4), 967–983. 10.1113/JP271830

Janak, P. H., & Tye, K. M. (2015). From circuits to behaviour in the amygdala. Nature, 517(7534), 284–292. 10.1038/nature14188

Kampa, B. M., Letzkus, J. J., & Stuart, G. J. (2006). Requirement of dendritic calcium spikes for induction of spike-timing-dependent synaptic plasticity. The Journal of Physiology, 574(1), 283–290. 10.1113/jphysiol.2006.111062

Koek, L. A., Sanderson, T. M., Georgiou, J., & Collingridge, G. L. (2024). The role of calcium stores in long-term potentiation and synaptic tagging and capture in mouse hippocampus. Philosophical Transactions of the Royal Society B: Biological Sciences, 379(1906). 10.1098/rstb.2023.0241

Kofler, J., Hattori, K., Sawada, M., DeVries, A. C., Martin, L. J., Hurn, P. D., & Traystman, R. J. (2004). Histopathological and behavioral characterization of a novel model of cardiac arrest and cardiopulmonary resuscitation in mice. J Neurosci Methods, 136(1), 33–44. 10.1016/j.jneumeth.2003.12.024

Krabbe, S., Grundemann, J., & Luthi, A. (2018). Amygdala Inhibitory Circuits Regulate Associative Fear Conditioning. Biol Psychiatry, 83(10), 800–809. 10.1016/j.biopsych.2017.10.006

Kwak, S., & Weiss, J. H. (2006). Calcium-permeable AMPA channels in neurodegenerative disease and ischemia. Curr Opin Neurobiol, 16(3), 281–287. 10.1016/j.conb.2006.05.004

Larson, J., & Munkacsy, E. (2015). Theta-burst LTP. Brain Res, 1621, 38–50. 10.1016/j.brainres.2014.10.034

LeDoux, J. (2007). The amygdala. Curr Biol, 17(20), R868–874. 10.1016/j.cub.2007.08.005

Li, X.-H., Miao, H.-H., & Zhuo, M. (2019). NMDA Receptor Dependent Long-term Potentiation in Chronic Pain. Neurochemical Research, 44(3), 531–538. 10.1007/s11064-018-2614-8

Luchkina, N. V., & Bolshakov, V. Y. (2019). Mechanisms of fear learning and extinction: synaptic plasticity–fear memory connection. Psychopharmacology, 236(1), 163–182. 10.1007/s00213-018-5104-4

Mahan, A. L., & Ressler, K. J. (2012). Fear conditioning, synaptic plasticity and the amygdala: implications for posttraumatic stress disorder. Trends Neurosci, 35(1), 24–35. 10.1016/j.tins.2011.06.007

Malenka, R. C. (1991). The role of postsynaptic calcium in the induction of long-term potentiation. Mol Neurobiol, 5(2-4), 289–295. 10.1007/BF02935552

Maren, S. (2005). Synaptic Mechanisms of Associative Memory in the Amygdala. Neuron, 47(6), 783–786. 10.1016/j.neuron.2005.08.009

Maren, S. (2017). Synapse-Specific Encoding of Fear Memory in the Amygdala. Neuron, 95(5), 988–990. 10.1016/j.neuron.2017.08.020

McDaid, J., Mustaly-Kalimi, S., & Stutzmann, G. E. (2020). Ca(2+) Dyshomeostasis Disrupts Neuronal and Synaptic Function in Alzheimer’s Disease. Cells, 9(12). 10.3390/cells9122655

Meis, S., Endres, T., Munsch, T., & Lessmann, V. (2018). The Relation Between Long-Term Synaptic Plasticity at Glutamatergic Synapses in the Amygdala and Fear Learning in Adult Heterozygous BDNF-Knockout Mice. Cerebral Cortex, 28(4), 1195–1208. 10.1093/cercor/bhx032

Mitchell, J. R., Trettel, S. G., Li, A. J., Wasielewski, S., Huckleberry, K. A., Fanikos, M., Golden, E., Laine, M. A., & Shansky, R. M. (2022). Darting across space and time: parametric modulators of sex-biased conditioned fear responses. Learn Mem, 29(7), 171–180. 10.1101/lm.053587.122

Naber, D., & Bullinger, M. (2018). Psychiatric sequelae of cardiac arrest. Dialogues in Clinical Neuroscience, 20(1), 73–77. 10.31887/DCNS.2018.20.1/dnaber

Nanou, E., & Catterall, W. A. (2018). Calcium Channels, Synaptic Plasticity, and Neuropsychiatric Disease. Neuron, 98(3), 466–481. 10.1016/j.neuron.2018.03.017

Noh, K.-M., Yokota, H., Mashiko, T., Castillo, P. E., Zukin, R. S., & Bennett, M. V. L. (2005). Blockade of calcium-permeable AMPA receptors protects hippocampal neurons against global ischemia-induced death. Proceedings of the National Academy of Sciences, 102(34), 12230–12235. 10.1073/pnas.0505408102

Oh, M. M., Oliveira, F. A., Waters, J., & Disterhoft, J. F. (2013). Altered Calcium Metabolism in Aging CA1 Hippocampal Pyramidal Neurons. The Journal of Neuroscience, 33(18), 7905–7911. 10.1523/jneurosci.5457-12.2013

Orfila, J. E., McKinnon, N., Moreno, M., Deng, G., Chalmers, N., Dietz, R. M., Herson, P. S., & Quillinan, N. (2018). Cardiac Arrest Induces Ischemic Long-Term Potentiation of Hippocampal CA1 Neurons That Occludes Physiological Long-Term Potentiation. Neural Plast, 2018, 9275239. 10.1155/2018/9275239

Peterlin, B. L., Nijjar, S. S., & Tietjen, G. E. (2011). Post-traumatic stress disorder and migraine: epidemiology, sex differences, and potential mechanisms. Headache, 51(6), 860–868. 10.1111/j.1526-4610.2011.01907.x

Phillips, R. G., & LeDoux, J. E. (1992). Differential contribution of amygdala and hippocampus to cued and contextual fear conditioning. Behav Neurosci, 106(2), 274–285. 10.1037//0735-7044.106.2.274

Phillips, R. G., & Ledoux, J. E. (1994). Lesions of the dorsal hippocampal formation interfere with background but not foreground contextual fear conditioning. Learning & Memory, 1(1), 34–44. 10.1101/lm.1.1.34

Pittig, A., Treanor, M., Lebeau, R. T., & Craske, M. G. (2018). The role of associative fear and avoidance learning in anxiety disorders: Gaps and directions for future research. Neuroscience & Biobehavioral Reviews, 88, 117–140. 10.1016/j.neubiorev.2018.03.015

Poon, C.-S., & Schmid, S. (2012). Nonassociative Learning. In Encyclopedia of the Sciences of Learning (pp. 2475–2477). 10.1007/978-1-4419-1428-6_1849

Quillinan, N., Grewal, H., Deng, G., Shimizu, K., Yonchek, J. C., Strnad, F., Traystman, R. J., & Herson, P. S. (2015). Region-specific role for GluN2B-containing NMDA receptors in injury to Purkinje cells and CA1 neurons following global cerebral ischemia. Neuroscience, 284, 555–565. 10.1016/j.neuroscience.2014.10.033

Rau, A. R., Chappell, A. M., Butler, T. R., Ariwodola, O. J., & Weiner, J. L. (2015a). Increased Basolateral Amygdala Pyramidal Cell Excitability May Contribute to the Anxiogenic Phenotype Induced by Chronic Early-Life Stress. J Neurosci, 35(26), 9730–9740. 10.1523/JNEUROSCI.0384-15.2015

Rau, A. R., Chappell, A. M., Butler, T. R., Ariwodola, O. J., & Weiner, J. L. (2015b). Increased Basolateral Amygdala Pyramidal Cell Excitability May Contribute to the Anxiogenic Phenotype Induced by Chronic Early-Life Stress. Journal of Neuroscience, 35(26), 9730–9740. 10.1523/jneurosci.0384-15.2015

Raybuck, J. D., & Lattal, K. M. (2011). Double dissociation of amygdala and hippocampal contributions to trace and delay fear conditioning. PLoS One, 6(1), e15982. 10.1371/journal.pone.0015982

Ressler, K. J. (2010). Amygdala activity, fear, and anxiety: modulation by stress. Biol Psychiatry, 67(12), 1117–1119. 10.1016/j.biopsych.2010.04.027

Rogan, M. T., Stäubli, U. V., & Ledoux, J. E. (1997). Fear conditioning induces associative long-term potentiation in the amygdala. Nature, 390(6660), 604–607. 10.1038/37601

Rosen, J. B., & Schulkin, J. (2022). Hyperexcitability: From Normal Fear to Pathological Anxiety and Trauma. Frontiers in Systems Neuroscience, 16. 10.3389/fnsys.2022.727054

Rumian, N. L., Chalmers, N. E., Tullis, J. E., Herson, P. S., & Bayer, K. U. (2021). CaMKIIalpha knockout protects from ischemic neuronal cell death after resuscitation from cardiac arrest. Brain Res, 1773, 147699. 10.1016/j.brainres.2021.147699

Sabatini, B. L., Oertner, T. G., & Svoboda, K. (2002). The Life Cycle of Ca2+ Ions in Dendritic Spines. Neuron, 33(3), 439–452. 10.1016/s0896-6273(02)00573-1

Sandler, V. M., & Barbara, J.-G. (1999). Calcium-Induced Calcium Release Contributes to Action Potential-Evoked Calcium Transients in Hippocampal CA1 Pyramidal Neurons. The Journal of Neuroscience, 19(11), 4325–4336. 10.1523/jneurosci.19-11-04325.1999

Seedat, S., Scott, K. M., Angermeyer, M. C., Berglund, P., Bromet, E. J., Brugha, T. S., Demyttenaere, K., De Girolamo, G., Haro, J. M., Jin, R., Karam, E. G., Kovess-Masfety, V., Levinson, D., Medina Mora, M. E., Ono, Y., Ormel, J., Pennell, B.-E., Posada-Villa, J., Sampson, N. A., … Kessler, R. C. (2009). Cross-National Associations Between Gender and Mental Disorders in the World Health Organization World Mental Health Surveys. Archives of General Psychiatry, 66(7), 785. 10.1001/archgenpsychiatry.2009.36

Seibenhener, M. L., & Wooten, M. C. (2015). Use of the Open Field Maze to Measure Locomotor and Anxiety-like Behavior in Mice. Journal of Visualized Experiments(96). 10.3791/52434

Siegel, C., Turtzo, C., & McCullough, L. D. (2010). Sex differences in cerebral ischemia: Possible molecular mechanisms. Journal of Neuroscience Research, 88(13), 2765–2774. 10.1002/jnr.22406

Simons, S. B., Escobedo, Y., Yasuda, R., & Dudek, S. M. (2009). Regional differences in hippocampal calcium handling provide a cellular mechanism for limiting plasticity. Proceedings of the National Academy of Sciences, 106(33), 14080–14084. 10.1073/pnas.0904775106

Smith, D. R., Gallagher, M., & Stanton, M. E. (2007). Genetic background differences and nonassociative effects in mouse trace fear conditioning. Learn Mem, 14(9), 597–605. 10.1101/lm.614807

Sun, Y., Gooch, H., & Sah, P. (2020). Fear conditioning and the basolateral amygdala. F1000Research, 9, 53. 10.12688/f1000research.21201.1

Sun, Y., Gooch, H., & Sah, P. (2020). Fear conditioning and the basolateral amygdala. F1000Res, 9. 10.12688/f1000research.21201.1

Tedoldi, A., Ludwig, P., Fulgenzi, G., Takeshima, H., Pedarzani, P., & Stocker, M. (2020). Calcium-induced calcium release and type 3 ryanodine receptors modulate the slow afterhyperpolarising current, sIAHP, and its potentiation in hippocampal pyramidal neurons. PLoS One, 15(6), e0230465. 10.1371/journal.pone.0230465

Theis, A.-K., Rózsa, B., Katona, G., Schmitz, D., & Johenning, F. W. (2018). Voltage Gated Calcium Channel Activation by Backpropagating Action Potentials Downregulates NMDAR Function. Frontiers in Cellular Neuroscience, 12. 10.3389/fncel.2018.00109

Tsvetkov, E., Carlezon, W. A., Benes, F. M., Kandel, E. R., & Bolshakov, V. Y. (2002). Fear conditioning occludes LTP-induced presynaptic enhancement of synaptic transmission in the cortical pathway to the lateral amygdala. Neuron, 34(2), 289–300. 10.1016/s0896-6273(02)00645-1

Turlova, E., Wong, R., Xu, B., Li, F., Du, L., Habbous, S., Horgen, F. D., Fleig, A., Feng, Z. P., & Sun, H. S. (2021). TRPM7 Mediates Neuronal Cell Death Upstream of Calcium/Calmodulin-Dependent Protein Kinase II and Calcineurin Mechanism in Neonatal Hypoxic-Ischemic Brain Injury. Transl Stroke Res, 12(1), 164–184. 10.1007/s12975-020-00810-3

Valdés-Undurraga, I., Lobos, P., Sánchez-Robledo, V., Arias-Cavieres, A., Sanmartín, C. D., Barrientos, G., More, J., Muñoz, P., Paula-Lima, A. C., Hidalgo, C., & Adasme, T. (2023). Long-term potentiation and spatial memory training stimulate the hippocampal expression of RyR2 calcium release channels. Frontiers in Cellular Neuroscience, 17. 10.3389/fncel.2023.1132121

Wickens, M. M., Bangasser, D. A., & Briand, L. A. (2018). Sex Differences in Psychiatric Disease: A Focus on the Glutamate System. Front Mol Neurosci, 11, 197. 10.3389/fnmol.2018.00197

Yan, S., Gan, Y., Jiang, N., Wang, R., Chen, Y., Luo, Z., Zong, Q., Chen, S., & Lv, C. (2020). The global survival rate among adult out-of-hospital cardiac arrest patients who received cardiopulmonary resuscitation: a systematic review and meta-analysis. Critical Care, 24(1). 10.1186/s13054-020-2773-2

Yang, S.-N., Tang, Y.-G., & Zucker, R. S. (1999). Selective Induction of LTP and LTD by Postsynaptic [Ca^2+^] _i_ Elevation. Journal of Neurophysiology, 81(2), 781–787. 10.1152/jn.1999.81.2.781

Yuste, R., Majewska, A., Cash, S. S., & Denk, W. (1999). Mechanisms of Calcium Influx into Hippocampal Spines: Heterogeneity among Spines, Coincidence Detection by NMDA Receptors, and Optical Quantal Analysis. The Journal of Neuroscience, 19(6), 1976–1987. 10.1523/jneurosci.19-06-01976.1999

