## Supplementary Figures for "Enhanced Calcium Signaling in BLA Pyramidal Neurons Underlies a Sex and Circuit Specific Amygdala Dysfunction After Global Cerebral Ischemia"

### Supplemental Figures:

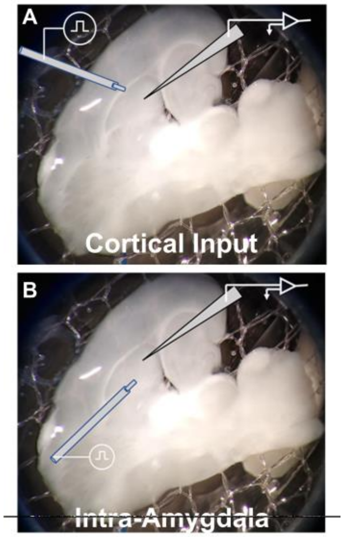


**Supplemental Figure 1.** To record fEPSPs in the cortical input, the stimulator and recording electrode were placed as depicted in (A). To record fEPSPs in the intra amygdala pathway, the stimulator and recording electrode were placed as depicted in (B).

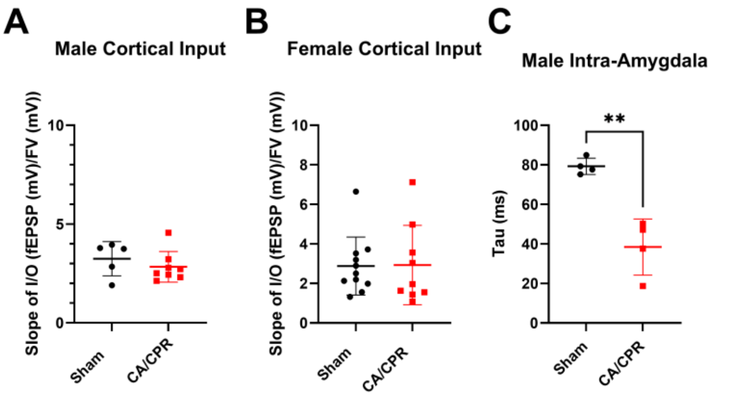


**Supplemental Figure 2: Quantification and comparison of Input/Output curves.** In recordings of the cortical input to the BLA, I/O responses were fit with a linear curve and slope values compared in male (A) and female (B) animals. In recordings of the intra-amygdala circuit, I/O responses were fit with an exponential curve and Tau values were compared in male animals (C). Students T-test. ** = $p < 0.01$

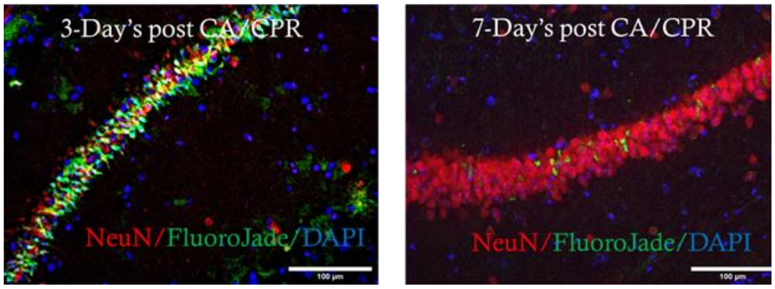


**Supplemental Figure 3: Neurodegeneration in the CA1 region of the hippocampus.** Representative images depicting profound neurodegeneration (green) in the hippocampus 3-days after CA/CPR (A), which subsides by 7-days post-CA/CPR (B).

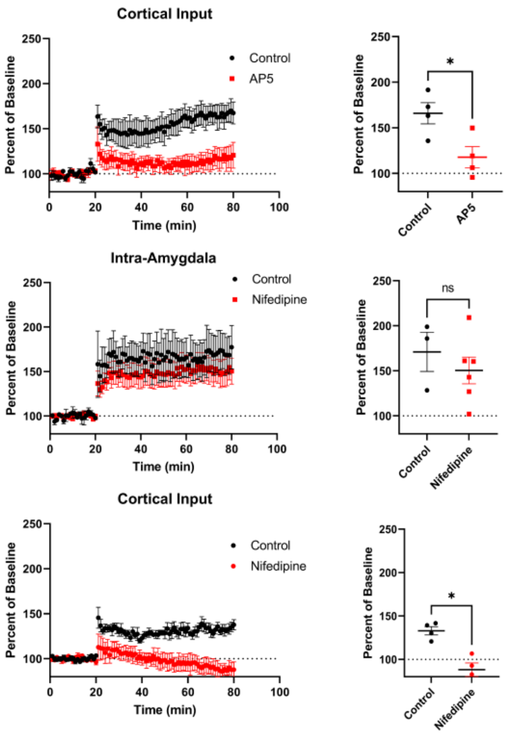


**Supplemental Figure 4: Pharmacological characterization of cortical input LTP and intra-amygdala LTP.** Inhibition of NMDA receptors with AP5 in the cortical input to the BLA during LTP recordings (A). Inhibition of LTCCs with Nifedipine while recording LTP of the intra-amygdala circuit (B). Inhibition of LTCCs with Nifedipine while recording LTP of the cortical input to the BLA (C). Students T-test. * = $p < 0.05$.
